# Dentate gyrus population activity during immobility supports formation of precise memories

**DOI:** 10.1101/2020.03.05.978320

**Authors:** Martin Pofahl, Negar Nikbakht, André N. Haubrich, Theresa Nguyen, Nicola Masala, Oliver Braganza, Jakob H. Macke, Laura A. Ewell, Kurtulus Golcuk, Heinz Beck

## Abstract

The hippocampal dentate gyrus is an important relay conveying sensory information from the entorhinal cortex to the hippocampus proper. During exploration, the dentate gyrus has been proposed to act as a pattern separator. However, the dentate gyrus also shows structured activity during immobility and sleep. The properties of these activity patterns at cellular resolution, and their role in hippocampal-dependent memory processes have remained unclear. Using dual-color in-vivo two-photon Ca^2+^ imaging, we show that in immobile mice dentate granule cells generate sparse, synchronized activity patterns associated with entorhinal cortex activation. These population events are structured and modified by changes in the environment; and they incorporate place- and speed cells. Importantly, they recapitulate population patterns evoked during self-motion. Using optogenetic inhibition during immobility, we show that granule cell activity during immobility is required to form dentate gyrus-dependent spatial memories. These data suggest that memory formation is supported by dentate gyrus replay of population codes of the current environment.

## Introduction

The dentate gyrus receives polymodal sensory information from the entorhinal cortex, and relays it into the hippocampal network. The most prevalent view of the dentate gyrus input-output transformation in this circuit is that it acts as a pattern separator. This capability requires the animal to generate dissimilar neuronal representations from overlapping input states that represent similar but not identical environments (***Cayco-Gajic and Silver, 2019***). Such an operation, termed pattern separation, has been ascribed to the hippocampal dentate gyrus in species ranging from rodents to humans (***Berron et al., 2016; Leutgeb et al., 2007; Sakon and Suzuki, 2019***). In the dentate gyrus, polysensory inputs are mapped onto a large number of granule cells which exhibit extremely sparse firing patterns, resulting in a high probability of non-overlapping output patterns (***Danielson et al., 2016; GoodSmith et al., 2017; Hainmueller and Bartos, 2018; Pilz et al., 2016; Senzai and Buzsáki, 2017; van Dijk and Fenton, 2018***). This concept has been influential in understanding dentate gyrus function when processing multimodal, current or ‘online’ sensory information during mobility and exploration.

However, the dentate gyrus is far from silent during immobility. It displays prominent electrographic activity patterns such as dentate spikes and sharp waves, which occur primarily during immobility or sleep (Bragin et al. 1995; Penttonen et al. 1997). In downstream hippocampal regions such as CA1, neuronal activity during immobility incorporates the replay of behaviorally relevant sequences during sharp wave ripples, a process important in memory consolidation (***Davidson et al., 2009; Diba and Buzsáki, 2007; Dupret et al., 2010; Foster and Wilson, 2006; Girardeau et al., 2009; Malvache et al., 2016; Skaggs and McNaughton, 1996; Wilson and McNaughton, 1994***). In the dentate gyrus, little detail is known about how granule cells are active during immobility and it is unknown whether activity during immobility reiterates behaviorally relevant information. Moreover, the role of dentate gyrus activity during immobility in memory formation is unclear.

Here, we have used dual-color two-photon in-vivo Ca^2+^ imaging to show that in immobile mice, the dentate gyrus exhibits frequent, sparse and synchronous population events that recapitulate activity patterns during locomotion at the population level. Moreover, we have tested the idea that activity during immobility is relevant for dentate-gyrus dependent spatial memory. We find that selective inhibition of granule cells only during immobility prevents the formation of dentate gyrus-dependent spatial memories.

## Results

### Sparse, structured dentate network events in immobile animals

We imaged the activity of >300 granule cells (GCs) using a Thy1-GCaMP6s mouse line (GP4.12Dkim/J, (***Dana et al., 2014***)). In addition, we monitored the activity of the major input system into the dentate gyrus, the medial perforant path (MPP). To this end, we expressed the red-shifted Ca^2+^ indicator jRGECO1a (***Dana et al., 2016***) in the medial entorhinal cortex using viral gene transfer (see Methods section, **Fig. 1A, B, Supplementary Fig. 1A**). To allow efficient excitation of both genetically encoded Ca^2+^ indicators, we established excitation with two pulsed laser sources at 940 and 1070 nm (see **Supplementary Fig. 1B**). The mice were placed under a two photon microscope and ran on a linear track (see **Supplementary Fig. 1C, Supplementary Movie 1**).

**Fig. 1:**
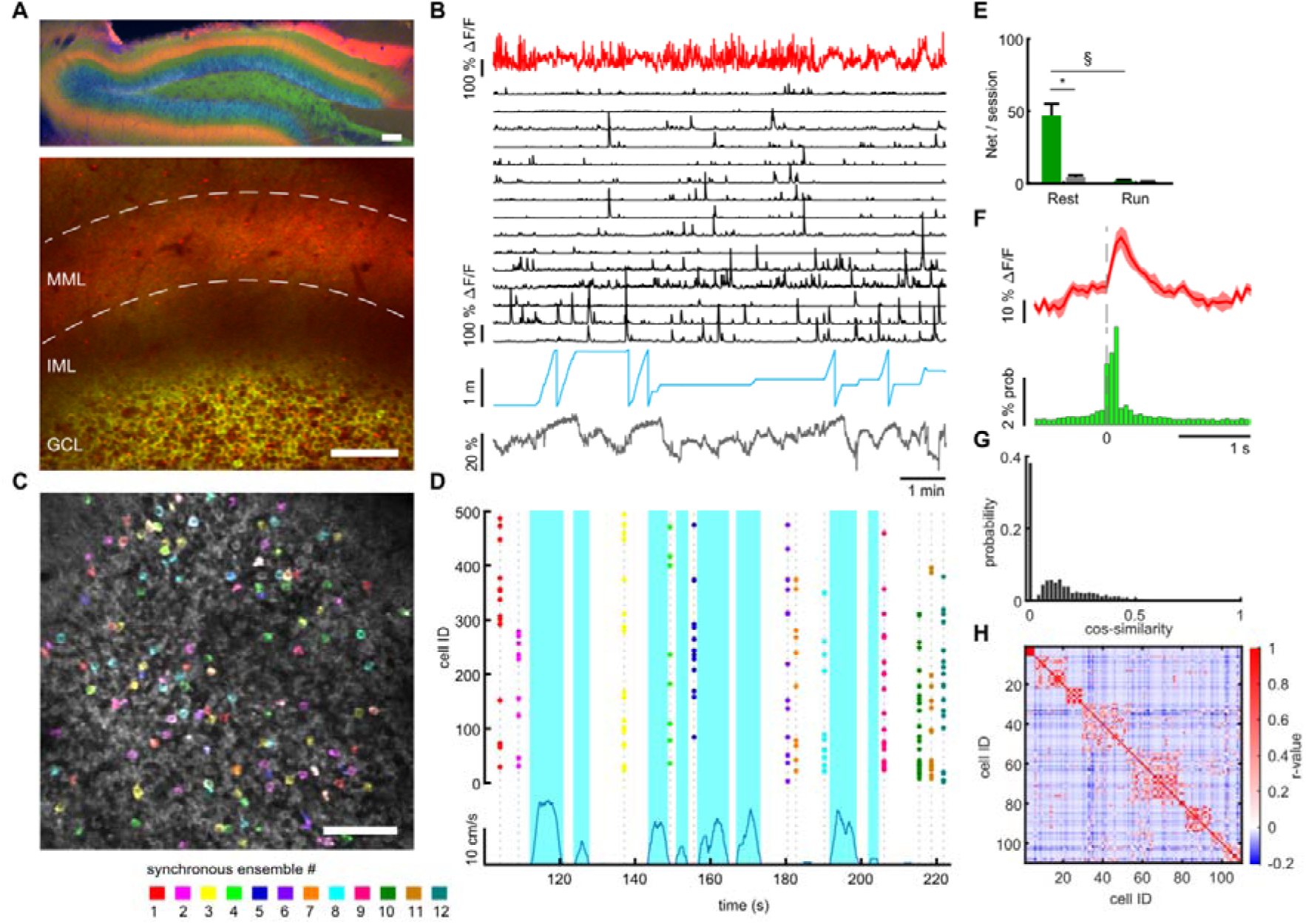
Synchronous MPP-driven dentate granule cell network events occur preferentially at rest. **A**, Expression of GCaMP6s in granule cells (Thy1-GCaMP mouse line, GP4.12Dkim/J). jRGECO was expressed in medial entorhinal cortex neurons using rAAV mediated gene transfer, and is visible in the middle molecular layer (MML) corresponding to the medial perforant path (MPP). Upper panel: Post hoc analysis in 70 µm fixed slice. Nuclei stained with DAPI (blue). Lower panel: Imaging plane for simultaneous recording of MPP bulk and individual GC activity. Scale bar 100 µm. **B**, Data of representative recording session. Bulk fluorescence signal of MPP fibers (red), extracted fluorescence signals from a subset of individual GCs (black), mouse position on linear track (blue) and diameter of mouse pupil (black). **C**, Participation of granule cells in synchronous network events. Representative field of view with highlighted simultaneously active GCs. Cells active during an individual network event are depicted in the same individual color. A subset of neurons is active in multiple network events, recognizable as white color. **D**, Raster plot of network events. Dashed lines mark network events, corresponding to simultaneous activity of > four cells. Participating cells are highlighted according to color scheme from panel C. Running speed is depicted (blue) to distinguish running and resting periods. **E**, Network events occurred mainly during immobility. Mean number of network events per recording session during running (light green) and resting (dark green). Grey bars depict shuffled data for each condition (n=9 animals). **F**, Average MPP activity (red) and probability of granule cells activity, both aligned to the time point of network events (n=3 animals, 538 granule cells). **G**, Similarity between network events. Similarity of population vectors computed for individual network events. Comparisons were carried out between all possible pairwise combinations of vectors and quantified using cosine similarity. **H**, Network activity contains ensembles that are repetitively active in a correlated fashion. Pairwise Pearson’s coefficients of granule cell activity during network events. Cells were sorted using hierarchical clustering to reveal highly correlated ensembles. Representative example.

As previously described, the firing of GCs was generally sparse (***Danielson et al., 2016; Hainmueller and Bartos, 2018; Neunuebel and Knierim, 2012; Pilz et al., 2016***), both when animals were immobile and running on a textured belt without additional cues (mean event frequency 1.38±0.19 events/min and 0.97±0.2 events/min, respectively, n=9 mice, **Fig. 1B, Supplementary Fig. 2C-F**). Despite the sparse activity of granule cells, we observed synchronized activity patterns. To rigorously define such events, we used an algorithm that detects synchronized network events within a 200 ms time window (see Methods). Such synchronous network events could readily be observed in the dentate gyrus (**Fig. 1C, D**, network events depicted in different colors, see corresponding **Supplementary Movie 2**). Network events were sparse, incorporating only 5.7±0.09 % of the active GC population. Notably, network events occurred mainly during immobility and were much less prevalent during running (**Fig. 1D, E**). Shuffling analysis (see Methods) confirmed that network events do not arise by chance (**Fig 1E**, grey bars correspond to shuffled data, ANOVA F_(3,25)_=30.12, p=4*10^−14^, Bonferroni post-test resting vs. shuffled p=2.2*10^−10^ indicated with asterisk, running vs. shuffled p=1). Simultaneous imaging of MPP and GC activity showed that network events were strongly correlated with MPP activity increases (**Fig. 1F**).

Moreover, cross-correlation revealed that during immobility, the increases in MPP activity were associated with peaks in average GC activity levels (**Supplementary Fig. 2F, J, K**). Both signals were significantly correlated in most sessions for periods of immobility (8/9 sessions, Granger causality test p<0.05), but not when the animals were running (8/9 sessions, Granger causality test p>0.05), suggesting that the MPP may contribute to driving network events.

We further characterized the participation of dentate granule cells in network events. We first asked in how far network events involve orthogonal cell populations. Indeed, while individual GCs can partake in multiple network events (see **Fig. 1B, C, Supplementary Movie 2**), we also observed network events that seemed completely distinct to others. To quantify how similar network events are to one another, we computed population vectors for each network event (see Methods). We then computed the cosine similarity as a measure of similarity between vectors representing individual network events, where identical patterns would have a cosine similarity of 1, and completely orthogonal patterns would exhibit a cosine similarity of 0. This analysis showed a large fraction of network event pairs that were completely orthogonal to one another, and significantly more than expected by chance (Wilcoxon test, p<0.005 comparison to shuffled data see **Supplementary Fig. 3A, B**). This is consistent with the capability to represent separate sets of information within network events (**Fig. 1G**).

Even though orthogonal network events were observed, we also observed a repeated activation of granule cells in multiple network events. To examine if specific sub-ensembles of granule cells are repeatedly recruited in network events, we performed a pairwise Pearson’s correlation of the activity of all cell pairs during all network events of a recording session (correlation coefficients depicted in the correlation matrix in **Supplementary Fig. 3C**). We then reordered the cells by hierarchical clustering (**Supplementary Fig. 3D**, more examples in **Supplementary Fig. 3F**). This analysis shows that subgroups of cells are strongly correlated within network event-related activity (**Fig. 1H**). We quantitatively defined clusters as those exhibiting a mean correlation coefficient within the cluster above chance level (**Supplementary Fig. 3D**, and Methods). Using this definition, the average cluster size was 6.7±0.4 cells per cluster. The repetitive nature of GC cluster activation during an entire session becomes clearly apparent when viewing cell activity during network events over an entire session, sorted by their participation in clusters (**Supplementary Fig. 3E**).

### Participation of place- and speed-coding granule cells in network events

We then identified the properties of the participating GCs, and asked if they encode specific spatial or locomotion-related information on a textured belt without cues as in Fig. 1. We first identified GCs that displayed significant position-related firing (**Fig. 2A** for representative polar plots of three GCs). Of the total number of granule cells, a small subgroup of GCs exhibited significant place coding (2.15%), with place fields distributed over the linear track (**Fig. 2B**). If the fraction of place-coding cells was calculated as a fraction of only GCs active during running as in Danielson et al. (2016), the fraction of significantly place-coding GCs was 23 % comparable to previously reported values (***Danielson et al., 2016***). Secondly, we identified a fraction of GCs (2.17 %) displaying a significant correlation of activity with running speed (**Fig. 2C-E**). This is in contrast to a previous study (***Danielson et al., 2016***), but consistent with data obtained in freely moving mice (***Stefanini et al., 2019***). We did not observe both speed coding and place coding in the same cells. Both place- and speed cells were similarly recruited into network events (55.42 % of place cells, 42.86 % of speed cells). These results show that MPP-driven network events in immobile animals incorporate neurons that code place- and speed related information during mobility.

**Fig. 2:**
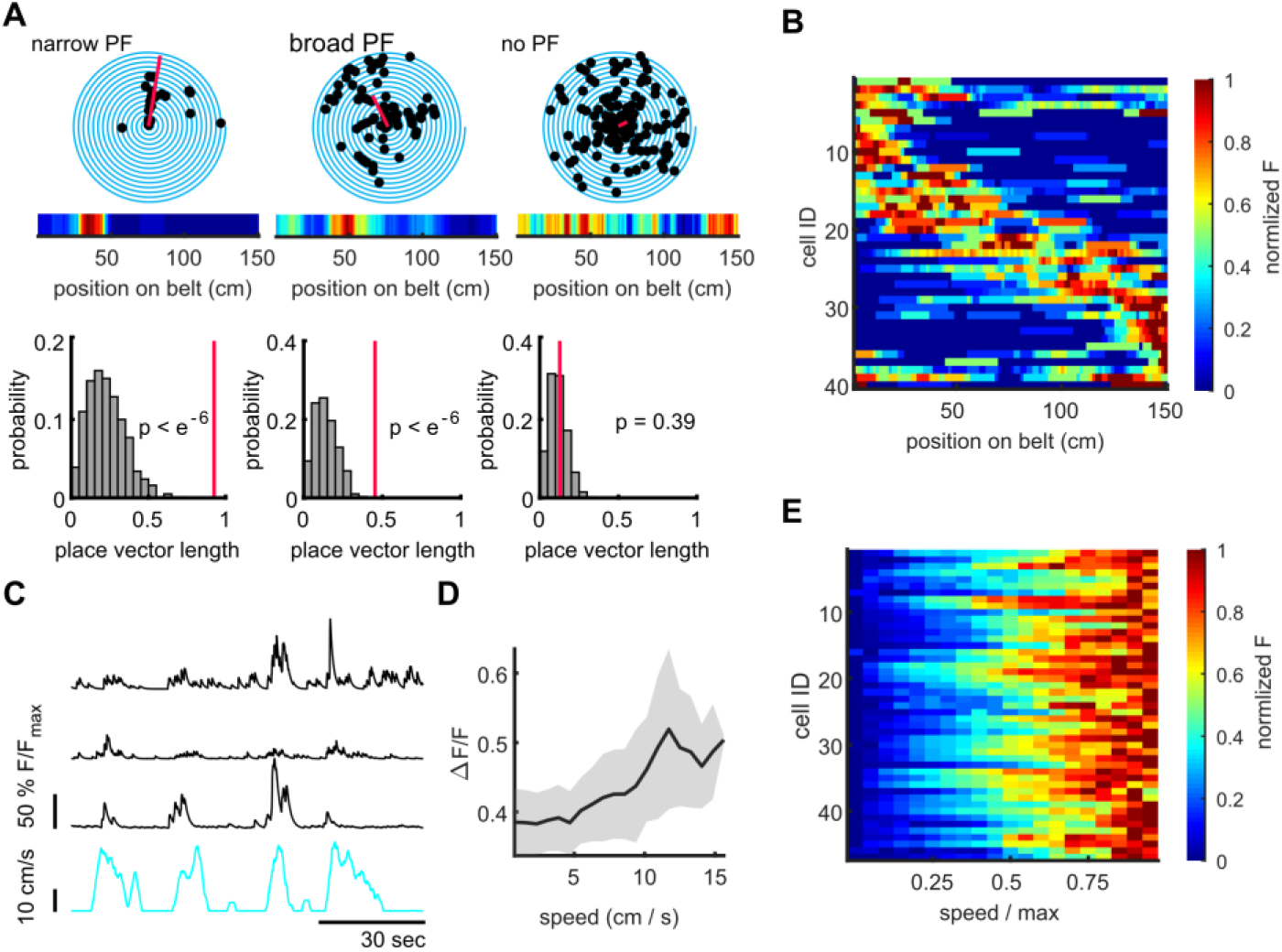
Characterization of dentate gyrus place and speed neurons. **A, B**, Place cells in the dentate gyrus. **A**, Representative polar plots of two significantly place-coding granule cells, and one without place coding. Place coding is depicted as spiral plot, where each 360° turn of the spiral represents a transition through the 1.5 m linear track. Detected events are shown as black dots. The red line represents the place vector. The corresponding heatmap of normalized fluorescence is shown in the inset. In lower panels, the distributions for place vector lengths generated from shuffled data (see Methods) are shown (grey histograms), the place vector for the individual cell is indicated as red line. **B**, Place field heatmaps of cells showing significant place preference. **C**, Representative examples of three significantly speed-coding neurons (black traces, running speed depicted in cyan). **D**, Speed-modulated mean fluorescence signal of a representative example cell. Gray area indicates standard error. **E**, Mean fluorescence signals of all significantly speed-modulated cells. Normalized fluorescence is color coded and running speed is normalized to every individual mouse maximum running speed.

### The properties of network events are altered when the environment changes

The incorporation of place cells suggests that information about the environment may be represented within network events. We therefore examined if the properties of network events are altered when the environment changes. To this end, we increased the density of sensory cues on the textured belt (cue-enriched condition, see methods, **Supplementary Fig. 1D-F**) in consecutive imaging sessions. In the cue-enriched condition, dentate gyrus network events were again observed predominantly during immobility (**Supplementary Fig. 4D**, statistics of GC activity in **Supplementary Fig. 4A-C**) and were similarly coupled to MPP activity (**Supplementary Fig. 4I-K**). Increasing the cue density did not significantly alter the network event frequency (2.39 ± 0.73 vs 3.63 ± 0.90 events/minute, respectively, n=9 mice, F_(1,30)_=4.32, p=0.34), or the fraction of GCs participating in network events in general (5.7±0.09 vs. 6.9±0.17 % of active GCs in baseline and cue-enriched conditions, respectively). However, the average size of individual network events, measured as the number of participating GCs, was significantly larger in the cue-enriched condition compared to the baseline condition (**Fig. 3A**, Kruskal-Wallis test, p=2*10^−37^), with individual GCs contributing more frequently to network events in the cue-rich condition (**Fig. 3B**, Kruskal-Wallis test, p=2*10^−26^). Fewer orthogonal networks were observed in the cue-rich condition, but this was not significantly different to the baseline condition (not shown, Kruskal-Wallis test, n.s. p=0.27).

**Fig. 3:**
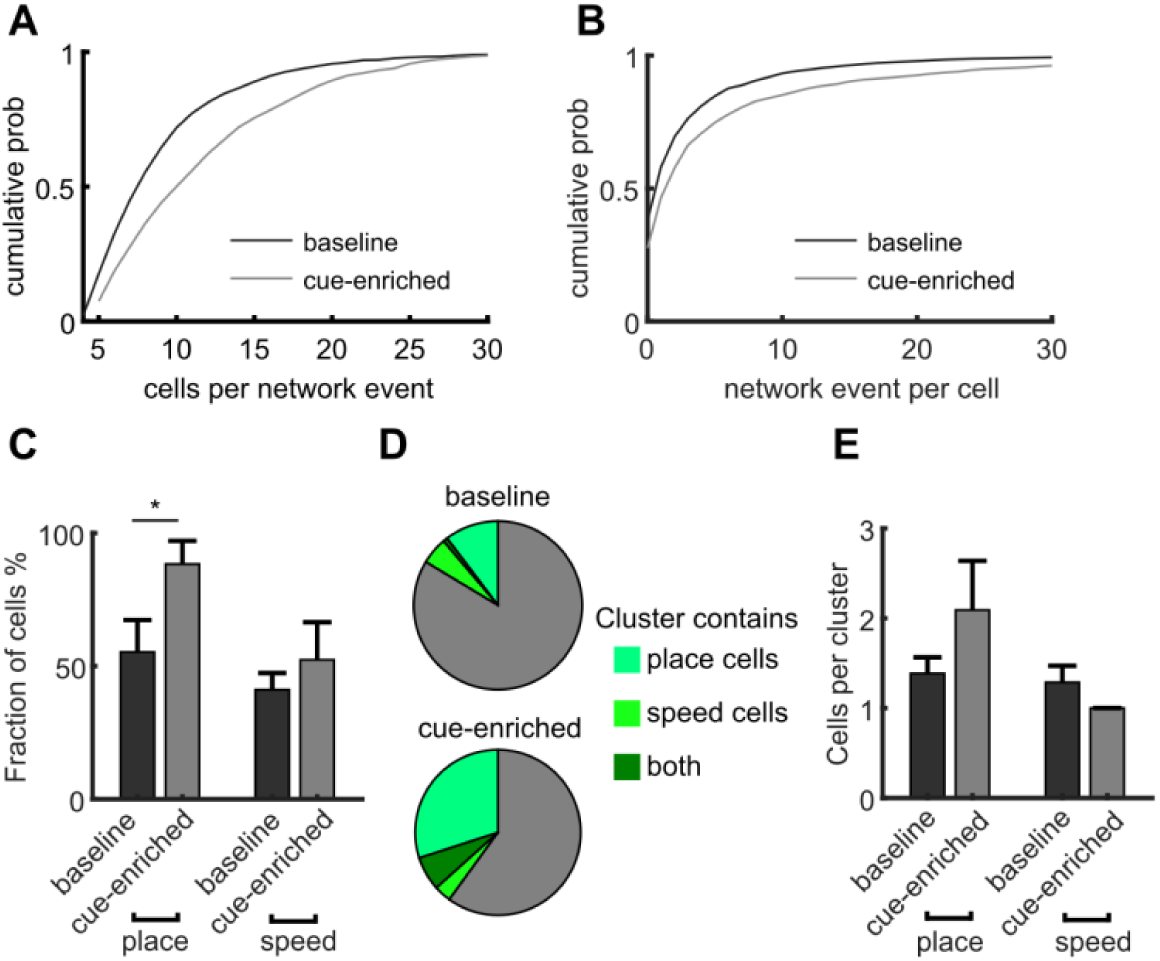
Increasing sensory cues causes enlargement of network events and increased incorporation of place cells. **A**, Network events involve more granule cells in cue-enriched environments. Cumulative probability of network event size (number of cells per network event) for baseline and cue enriched condition (dark and light grey lines, respectively). **B**, Cumulative probability of participation in multiple network events per cell for baseline and cue enriched condition (dark and light grey lines, respectively). **C**, Fraction of place and speed cells that participate in network events (total number of place/speed cells equals 100%). **D**, Relation of place and speed cells to correlated cell clusters (c.f. Fig. 1H). Fraction of the total number of clusters containing place cells (dark green), speed cells (light green), or both (intermediate green). Gray indicates clusters containing neither place nor speed cells. **E**, Mean number of place and speed cells per cluster, in baseline and cue enriched conditions. (n=9 animals)

We then examined if the participation of place and speed cells in network events is altered by increasing sensory cue density. Place cells were more commonly detected in the cue-enriched condition (4.66 vs. 2.15% of GCs), while there was no significant change in the proportion of speed cells (2.17 vs. 1.7 % of GCs in the baseline and cue-rich conditions, respectively, chi^2^ test regarding changes in the fraction of place and speed cells p=6*10^−4^, post-test: place cells baseline vs. cue-enriched p=1.7*10^−4^, speed cells baseline vs. cue-enriched p=0.33).

For place cells, the probability of being incorporated in network events was increased significantly (**Fig. 3C**, 55.42 vs. 88.46 % of place cells in baseline vs. cue-rich conditions), while this was not the case for speed cells (42.86 vs. 52.63 % of speed cells in baseline vs. cue-rich conditions, chi^2^ test regarding changes in the incorporation of place and speed cells in network events p=1*10^−4^, post-test: place cells baseline vs. cue-enriched p=3*10^−6^, indicated with asterisk in **Fig. 3C**, speed cells baseline vs. cue-enriched p=0.32). The properties of correlated cell clusters within network events did not change (cluster size comparison, Kruskal-Wallis test, p=0.16), but significantly more of the clusters contained place cells in the cue-rich condition (**Fig. 3D**, Chi^2^ test p=0.012, post-test comparison baseline vs. cue-enriched for place cells p=0.02, speed cells p=0.20), with the number of place or speed cells per cluster remaining unchanged (**Fig. 3E**). Thus, network events are responsive to changes in the environment, and incorporate more place-coding neurons into correlated activity patterns.

### Similarity of population activity patterns during locomotion to network events during immobility

The incorporation of place and speed cells in network events, as well as the fact that changing features of the environment modifies network event size and place cell participation is consistent with the idea that animals, when immobile, represent information about the environment in synchronous, sparse network events. Testing this idea is difficult, however, given that place cells are less prevalent in the dentate gyrus compared to other hippocampal subregions. It has been suggested that the dentate gyrus utilizes a population code (***Stefanini et al., 2019***), meaning that even though only few cells can be rigorously classified as place cells, many more neurons may encode relevant but partial information about the environment. We therefore developed an approach to assess similarity between running and resting activity in the dentate gyrus at the population level.

We analyzed population coding during both locomotion and immobility in DG using Principal Component Analysis (PCA) of the activity during individual imaging sessions. We visualized the neuronal state during locomotion captured by the first three components (**Fig. 4A, E, Supplementary Fig. 5B-C** for Independent Component Analysis, ICA, and Gaussian Process Factor Analysis, GPFA). We observed smooth, large trajectories with high variability reflecting movement along the linear track for some revolutions on the linear treadmill. Because these trajectories did not repeat across multiple rounds, we validated the PCA approaches by applying them to population activity in CA1, in which population codes representing space studied with PCA have been found to be repetitive and stable across different iterations of similar behavior (***Rubin et al., 2019***). Indeed, in CA1 trajectories were consistent and stable across multiple rounds, and related smoothly to the position on the linear track (**Supplementary Fig. 5D-F**, for PCA, GPFA, ICA, see Methods). After applying PCA to locomotor states in the dentate gyrus, we then also performed a PCA analysis of the population activity in the same region during network events (see Methods, **Fig. 4B**).

**Fig. 4:**
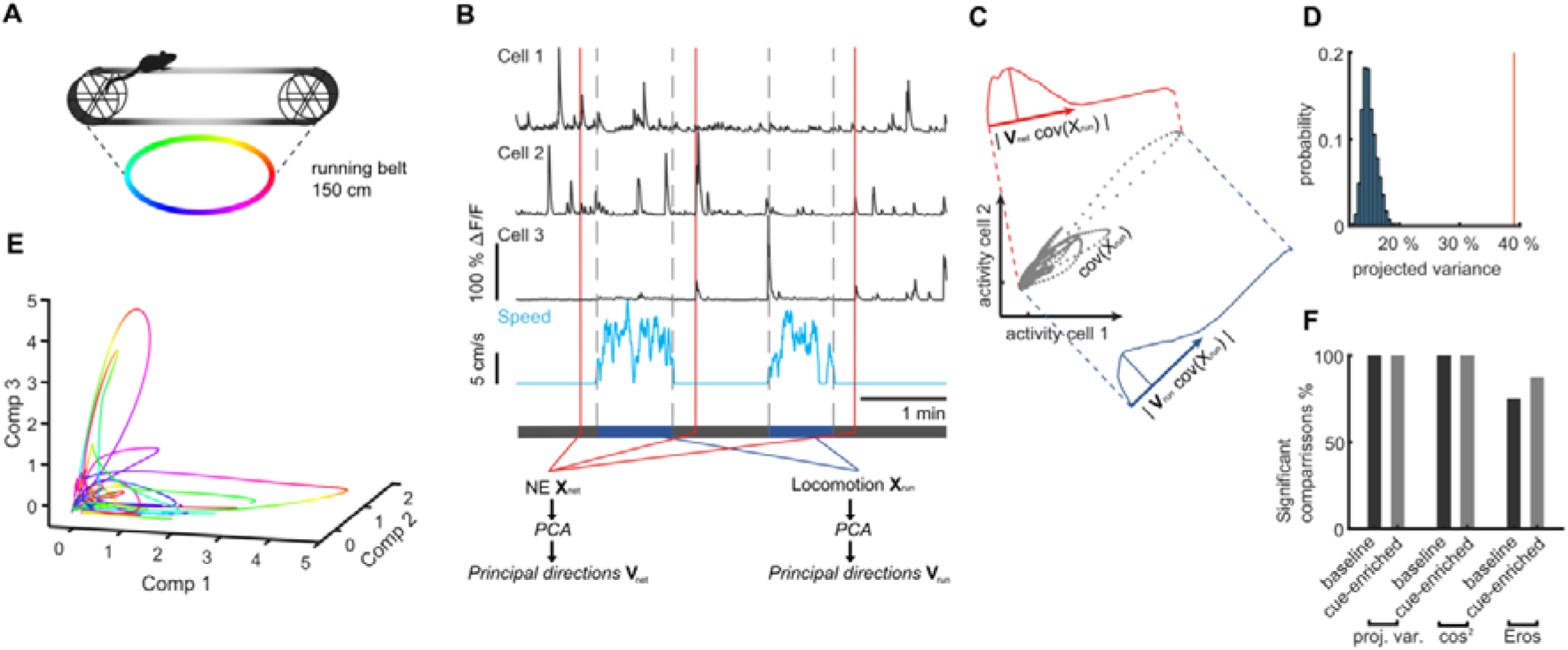
Similarity of activity patterns during network events to population patterns during locomotion. **A**, Color code for position on the linear track used in panels E. **B**, schematic of the procedure for comparing population activity during network events (NE) and locomotion. Population activity is represented by 3 cells (upper traces), recorded during running and quiet immobility (blue trace indicates speed). Time point of three network events is indicated schematically by red lines. Activity during network events (NE) was used to perform PCA, computing the transformation matrix V_net_. Similarly, PCA was performed on the neuronal population activity only from running periods (speed indicated in blue, bordered by vertical grey dashed lines), to generate the transformation matrix V_run_ representing the covarying activity during locomotion. **C**, Schematic description of the procedure for projecting co-variances of running activity into the PCA basis of network events (or shuffled data). Grey dots show covarying activity of two representative cells during running. The blue graph denotes the projection into the locomotion PCA-space using V_run_ and the width of the distribution shows the projected variance. The red graph shows the same information for the network space using V_net_. **D**, Individual example of shuffle analysis for a representative session, where the vertical red line indicates the projected variance explained normalized to original variance. The variance explained is larger than the shuffled distribution (blue), indicating that the population activity during locomotion and network events is more similar than expected by chance (i.e. for network activity without correlations). **E**, Trajectories during an individual representative session plotted in a three-dimensional coordinate system corresponding to the first three PCA components (Comp 1-3). **F**, Fraction of sessions in which comparisons of population activity were significant vs. chance level for the two cue conditions and all similarity measures (n = 9 mice, 1 session per condition, see Supplementary Fig. 7 for comparisons to shuffled datasets for all sessions).

In order to compare the two sets of PCAs representing population activity during running states and network events, respectively, we first used a vector-based similarity measure. Briefly, we projected the traces recorded during locomotion into the PCA-space representing activity during network events, and tested how much of their variance was captured by them. In this analysis, similarity between both population measures would result in a large fraction of explained variance (see **Fig. 4B, C**). To obtain the expected null distribution, we performed a shuffling analysis on the resting activity to eliminate inter-neural correlations (see Methods). The distributions from shuffled data were clearly distinct from the real data (red vertical line in **Fig. 4D, Supplementary Fig. 6** for comparisons to shuffled data for all sessions). The comparisons to shuffled data were significant in all sessions, both for baseline and cue-enriched conditions (**Fig. 4F**, leftmost bars), indicating that the activity during network events is more similar to activity during locomotion than expected by chance.

In addition to this measure of similarity, we used two further measures based on cosine similarity (***Krzanowski, 1979***) and EROS (***Yang, K., Shahabi, C., 2004***), testing them against shuffled datasets in the same manner (**Fig. 4F, Supplementary Fig. 6** for comparisons to shuffled data for all sessions). With these measures, significant comparisons to shuffled data were obtained with all (cosine similarity) or a majority (EROS) of sessions. All three measures thus show that the activity during network events is more similar than expected by chance to sensory-driven activity during locomotion.

### Inhibition of dentate granule cell activity during immobility disrupts pattern separation

Collectively, these data suggest that during immobility, GCs engage in structured ensemble activity that reiterates activity during running at the population level. This suggests that such activity might be important for the formation of hippocampal dependent spatial memories. We tested this hypothesis using an established memory task for spatial object pattern separation (OPS, (***van Goethem et al., 2018***)), in which DG-dependent spatial discrimination is assessed based on the differential exploration of two objects. Briefly, animals are first exposed to two objects in defined locations during an acquisition trial (5 min) and are then re-exposed following an intermediate period, with one of the objects slightly displaced. Increased exploration of the displaced object indicates that the animal has encoded the initial location and is able to discriminate the displaced object. In preliminary experiments, we tested 4 degrees of object displacement along a vertical axis (3-12 cm, **Supplementary Fig. 7C**). We then determined the extent to which spatial object pattern separation was dependent on the activity of the dentate gyrus. We expressed either halorhodopsin (eNpHR, (***Gradinaru et al., 2008***)), or eYFP (control group) selectively in dentate GCs using Prox1-Cre mice, which efficiently inhibited GC firing (**Supplementary Fig. 7E-J**), and bilaterally illuminated the dentate gyrus with two implanted light fibers during the OPS task (**Supplementary Fig. 7A, B**). We found that GC activity was most important for an intermediate degree of displacement (9 cm), while maximal displacement was no longer dependent on GC activity (**Supplementary Fig. 7D**).

We then used the intermediate degree of displacement in the OPS task to see if dentate gyrus activity during quiet immobility was necessary to acquire a memory of object location. We bilaterally inhibited GCs during periods of quiet immobility (running speed < 4 cm/s) only during the learning trial (**Fig. 5A, B**). This manipulation led to a complete loss of preference for the displaced object in the subsequent recall trials (**Fig. 5H**, for representative sessions, analysis of discrimination index in **I**, n=6, T-test with Welch correction p=0.0002 for comparison between eNpHR and eYFP group) whereas control mice displayed a clear preference for the displaced object (**Fig. 5G** for representative session, n=9).

**Fig. 5:**
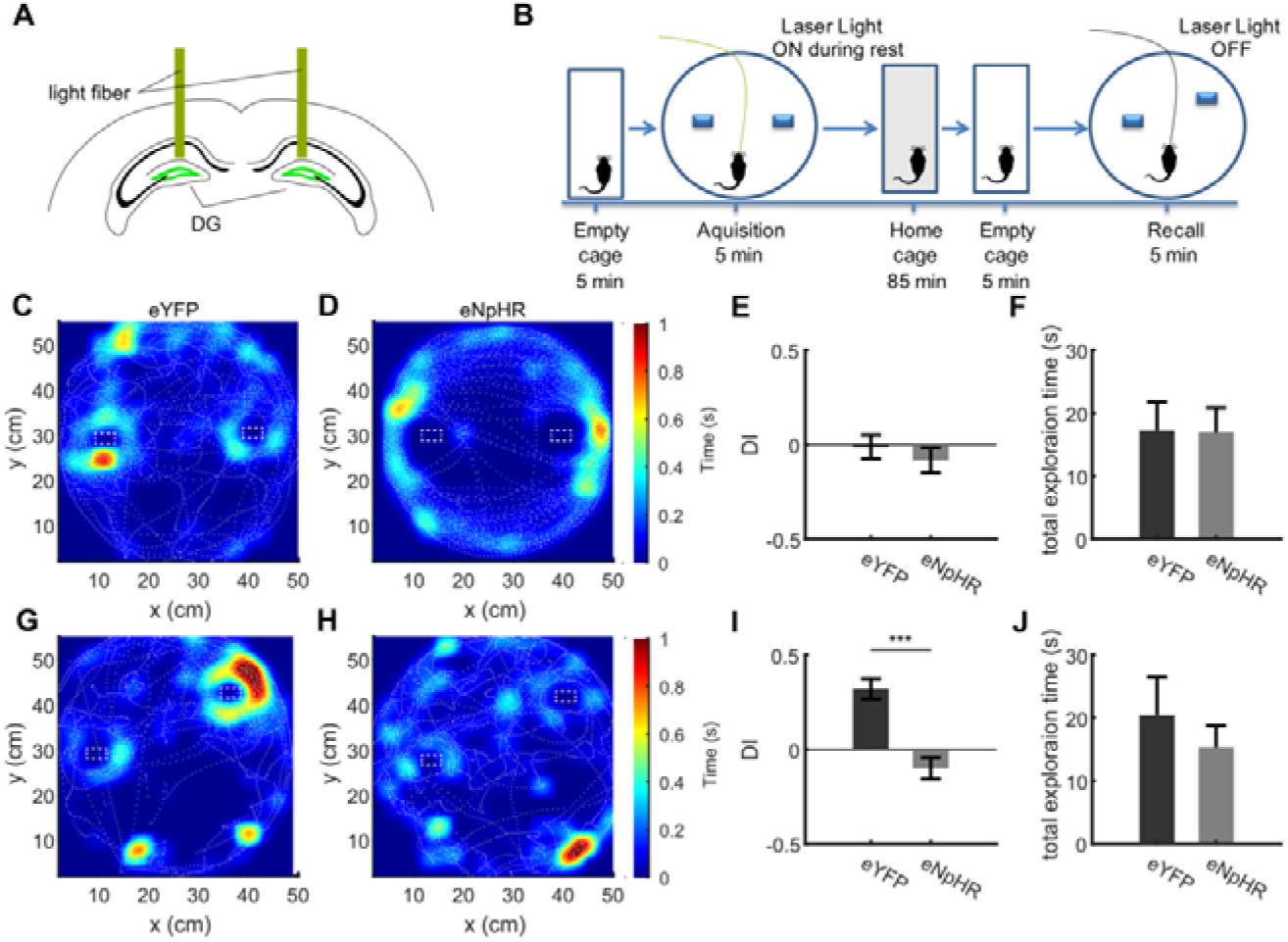
Inhibition of dentate granule cell activity during immobility prevents memory acquisition. **A**, Schematic of the bilateral optogenetic inhibition of the dentate gyrus granule cells expressing eNpHR. **B**, Schematic of the experimental procedure. In the acquisition phase, mice were familiarized with an arena containing two objects. Following an intermediate period of 90 minutes, the mice were placed in the same arena in which one object was moved slightly. **C, D**, Representative sessions from acquisition trials in control (eYFP) mice and mice expressing eNpHR in granule cells showing the tracking of the mouse center of mass (dashed white lines), as well as normalized occupancy within the arena. **E**, Discrimination index from the acquisition trial quantifying the specific exploration activity of the objects relative to one another (see Methods), with 0 values indicating equal exploration (see Methods). **F**, Total time spent exploring the objects in the eYFP and eNpHR groups during the acquisition trial. **G, H**, Representative sessions from recall trials depicted as shown in B, C. **I**, Discrimination index for recall trials, showing strong preference for the displaced object in the eYFP group, but not the eNpHR group if granule cell activity was inhibited during acquisition trials only during immobility. **J**, Total time spent exploring the objects in the eYFP and eNpHR groups during the recall trial (n=6 animals for eNpHR group, n=9 animals for eYFP group).

Carrying out the OPS task in mice expressing eNpHR without illumination yielded discrimination indices indistinguishable from the control group (not shown). For the three groups, ANOVA revealed a significant effect (F=8,515 p=0.0025), with Bonferroni post-tests showing that inhibition of GCs significantly reduces performance vs. the two control groups (eYFP vs. eNpHR illuminated p=0.0056, eYFP vs. eNpHR without illumination p>0.99, eNpHR with illumination vs. without illumination p=0.0059).

Because mice are also immobile while examining the objects, we also performed a set of experiments in which light-stimulation was only carried out during immobility, but excluding a 4 cm zone surrounding the objects (**Supplementary Fig. 8A**). This experiment yielded similar results, with a virtually complete loss of object discrimination during the recall trial (**Supplementary Fig. 7B-I**). This effect was specific to acquisition. GC inhibition only during immobility in the recall trial (**Supplementary Fig. 8J**) elicited no significant reduction in the recognition of the displaced object (**Supplementary Fig. 8K-L**, n=11 and 6 for eYFP and eNpHR groups, respectively, t-test with Welch correction n.s.). These data suggest that activity of GCs during rest is important to form memories that require discrimination of similar experiences.

## Discussion

The dentate gyrus has been implicated in pattern separation of sensory-driven activity patterns during experience, but is also active during immobility and sleep. The properties of these latter forms of activity at the cellular level and the role they play in behavior are largely unknown. The application of multiphoton in-vivo Ca^2+^ imaging allowed us to observe large-scale dentate gyrus dynamics at the cellular level and to detect a novel form of sparse, synchronized GC activity that occurs during immobility, termed dentate network events. These events were specifically modified by the environment, showed higher similarity than expected by chance to population activity occurring during locomotion, and were important for formation of memories requiring pattern separation. This suggests that sparse reiteration of locomotion-associated activity patterns is important for the formation of dentate gyrus-dependent memories.

For neuronal activity during resting states to support learning or memory consolidation concerning a particular environment, one general requirement would be that it recapitulates aspects of the population activity induced by exploration of the relevant environment (***Davidson et al., 2009; Diba and Buzsáki, 2007; Dupret et al., 2010; Foster and Wilson, 2006; Girardeau et al., 2009; Skaggs and McNaughton, 1996; Wilson and McNaughton, 1994***). We have used three different similarity measures to show that this is the case in the dentate gyrus at the population level.

In addition to this general requirement, two specific features of resting activity are consistent with the formation of precise memories that conserve the pattern separation capabilities of the dentate gyrus. Firstly, the activity patterns, although sparse, should be capable of generating orthogonal ensembles representing different features of the environment. Secondly, the activity should repetitively recruit specific subsets of dentate GCs capable of instructing the formation of CA3 attractors via Hebbian plasticity mechanisms. Indeed, we found that activity during network events is sparse, recruiting just ∼5-7% of the active GCs. Given that only ∼50% of GCs are active in head-fixed animals (***Danielson et al., 2016; Pilz et al., 2016***), recruitment of dentate GCs during network events is much sparser than in other forms of activity occurring during immobility or sleep. For instance, the fraction of CA1 neurons recruited during sharp wave ripple mediated replay of behaviorally relevant sequences (***Davidson et al., 2009; Diba and Buzsáki, 2007; Dupret et al., 2010; Foster and Wilson, 2006; Girardeau et al., 2009; Malvache et al., 2016; Skaggs and McNaughton, 1996; Wilson and McNaughton, 1994***) is much higher than the recruitment of GCs in network events. One consequence of the sparseness of network events is that they are predicted to recruit highly constrained CA3 ensembles, both because of the sparse excitatory connectivity of mossy fibers in CA3, and because of the properties of the powerful inhibitory circuits in the CA3 region (***Acsady et al., 1998; Neubrandt et al., 2017; Neubrandt et al., 2018***). This has been suggested to be important in the capability to store information in CA3, while conserving the pattern separation benefits of the dentate gyrus (***O’Reilly and McClelland, 1994; GoodSmith et al., 2019***).

We found that network events are structured, with subgroups of dentate GCs forming correlated subassemblies that are repeatedly recruited (see Supplementary Figure 3c). This finding is consistent with the idea that dentate GC activity recruits plasticity mechanisms to form sparse attractor-like representations in CA3 (***O’Reilly and McClelland, 1994***). A similar structure was also observed for awake hippocampal reactivations in the hippocampal CA1 region, and may serve similar plasticity mechanisms in downstream targets (***Malvache et al., 2016***). Thus, dentate activity during immobility may be important to instruct downstream ensembles to exhibit specific memory-related sequences. That the integrity of the dentate gyrus is important in determining behaviorally relevant firing patterns in CA3 has also been demonstrated by lesion experiments showing that activity of dentate GCs is necessary for increased SWRs and prospective goal-directed firing of CA3 neurons (***Sasaki et al., 2018***).

Our behavioral data are consistent with the hypothesis that dentate gyrus network events during immobility are important to generate dentate-dependent spatial memories. The OPS task requires storage of the initial object location with a high degree of precision that can be utilized later on for discriminating the translocated object. We show that optogenetically inhibiting GCs, even if this was done only during immobility remote from the explored objects, disrupted the capability to acquire such memories. This suggests that dentate network events rapidly and flexibly introduce information about the environment into the hippocampal network, in the time intervals interspersed between episodes of exploration. Consistent with this view of ‘real-time updating’, we observed increased incorporation of spatial information via place cell integration into network events upon the first experience of a cue-rich environment. Inhibition of the dentate gyrus during the recall phase did not significantly inhibit task performance, consistent with the idea that recall of precise location information is achieved via activation of memory-related attractors in downstream CA3 and/or CA1 regions. Indeed, behavioral analyses combined with selective lesions of dentate gyrus and CA3 have also suggested an interaction between CA3 and DG in supporting encoding but not retrieval processes in a spatial learning task (***Jerman et al., 2006***). Moreover, disrupting dentate spikes via electrical stimulation has been shown to disrupt acquisition of hippocampal-dependent trace eyeblink conditioning (Nokia et al. 2017).

How do network events correspond to the different types of activity that have been described in the dentate gyrus during immobility or sleep, namely dentate spikes and sharp waves (Bragin et al. 1995; Penttonen et al. 1997)? During dentate spikes, granule cells are discharged anterogradely by entorhinal input, while they are activated retrogradely by the CA3-mossy cell feedback pathway during sharp waves (Bragin et al. 1995; Penttonen et al. 1997). We show that dentate network events are associated with MPP activation, but would be cautious in designating these events dentate spikes in the absence of parallel in-vivo electrophysiology.

In addition to the formation of precise memories, activity in the dentate gyrus may be relevant for processes on more extended time scales, such as maintenance of established memories. For instance, pharmacogenetic inhibition of GCs induces loss of a hippocampal memory in trace eyeblink conditioning (***Madroñal et al., 2016***). Such longer time-scale coding may be mediated by processes that extend beyond local hippocampal computations. Along these lines, dentate gyrus activity during dentate spikes is associated with wide-spread increases in single-cell activity, gamma oscillations and intraregional gamma coherence (***Headley et al., 2017***). It is thus possible that the precise activation patterns we observe in dentate gyrus here are part of a more distributed, organized activity occurring in immobile animals.

In summary, we have described a novel form of synchronized, sparse network activity during immobility in DG, and have demonstrated that DG activity during immobility is required for the formation of precise spatial memories. Such DG network events may drive synchronous, restricted ensembles in CA3, creating multiple orthogonal, sparse attractor-like representations important for precise memories.

## Methods

### Animals and procedures

All experimental procedures were conducted in accordance to federal law of the state of North Rhine-Westphalia. We used 9-12 weeks old Thy1-GCaMP6 mouse line (GP4.12Dkim/J) mice for imaging experiments, which express GCaMP6s in most hippocampal neurons (***Dana et al., 2014***). For optogenetic inhibition of the dentate gyrus granule cells, we used heterozygous Prox1-Cre animals (Tg(Prox1-cre)SJ39Gsat/Mmucd) obtained from MMRRC UC Davis as cryopreserved sperm and rederived in the local facility.

### Virus injections and head fixation

Thy1-GCaMP6 mice were anesthetized with a combination of fentanyl/midazolam/medetomidine (0.05 / 5.0 / 0.5 mg/kg body weight i.p.) and head-fixed in a stereotactic frame. 30 min prior to induction of anesthesia, the animals were given a subcutaneous injection of ketoprofen (5 mg/kg body weight). Eyes were covered with eye-ointment (Bepanthen, Bayer) to prevent drying and body temperature was maintained at 37°C using a regulated heating plate (TCAT-2LV, Physitemp) and a rectal thermal probe. After removal of the head hair and superficial disinfection, the scalp was removed about 1 cm^2^ around the middle of the skull. The surface was locally anesthetized with a drop of 10% lidocaine and after 3-5 min residual soft tissue was removed from the skull bones with a scraper and 3% H_2_O_2_/NaCl solution. After complete drying, the cranial sutures were clearly visible and served as orientation for the determination of the drilling and injection sites. For virus injection, a hole was carefully drilled through the skull with a dental drill, avoiding excessive heating and injury to the meninges. Any minor bleeding was stopped with a sterile pad. The target site was located as the joint of Parietal, Interparietal and Occipital skull plates. Subsequently, the tip of a precision syringe (cannula size 34 G) was navigated stereotactically through the burrhole (30° towards vertical sagittal plane, 1.5 mm depth from skull surface) and virus particles (rAAV2/1-CaMKIIa-NES-jRGECO1a (***Dana et al., 2016***)) were slowly injected (total volume 250 nl, 50 nl / min) in the medial entorhinal cortex. Correct injection site in the medial entorhinal cortex was verified in all cases by confined expression of jRGECO1a in the middle molecular layer of the dentate gyrus (**Fig. 1a**). To prevent reflux of the injected fluid, the cannula was retained for 5 minutes at the injection site. Optibond (Optibond™ 3FL; two component, 48% filled dental adhesive, bottle kit; Kerr; FL, USA) was then applied thinly to the skull to aid adhesion of dental cement. Subsequently, a flat custom made head post ring was applied with the aid of dental cement (Tetric Evoflow), the borehole was closed and the surrounding skin adapted with tissue glue, also closing the borehole and adapting the surrounding skin with tissue glue. At the end of the surgery, anesthesia was terminated by i.p. injection of antagonists (naloxone/flumazenil/atipamezole, 1.2 / 0.5 / 2.5 mg / kg body weight). Postoperative analgesia was carried out over 3 days with 1 × daily ketoprofen (5 mg/kg body weight, s.c.).

### Window implantation procedure

Cranial window surgery was performed to allow imaging from the hippocampal dentate gyrus. 30 minutes before induction of anesthesia, the analgesis buprenorphine was administered for analgesia (0.05 mg/kg body weight) and dexamethasone (0.1 mg/20 g body weight) was given to inhibit inflammation. Mice were anesthetized with 3-4% isoflurane in an oxygen/air mixture (25/75%) and then placed in a stereotactic frame. Eyes were covered with eye-ointment (Bepanthen, Bayer) to prevent drying and body temperature was maintained at 37°C using a regulated heating plate (TCAT-2LV, Physitemp) and a rectal thermal probe. The further anesthesia was carried out via a mask with a reduced isoflurane dose of 1-2% at a gas flow of about 0.5 l / minute. A circular craniotomy (Ø 3 mm) was opened above the right hemisphere hippocampus using a dental drill. Cortical and CA1 tissue was aspirated using a blunted 27-gauge needle until the blood vessels above the dentate gyrus became visible. A custom made cone-shaped silicon inset (Upper diameter 3 mm, lower diameter 1.5 mm, length 2.3 mm, RTV 615, Movimentive) attached to by a cover glass (Ø 5 mm, thickness 0.17 mm) was inserted and fixed with dental cement. This special window design allowed easy implantation and maintenance and minimized the amount of aspirated tissue. Further the geometry was optimal for conserving the numerical aperture of the objective (**Supplementary Fig. 9**). Postoperative care included analgesia by administering buprenorphine twice daily (0.05 mg/kg body weight) and ketoprofen once daily (5 mg/kg body weight s.c.) on the three consecutive days after surgery. Animals were carefully monitored twice daily on the following 3 days, and recovered from surgery within 24-48 hours, showing normal activity and no signs of pain. The preparation of CA1 imaging windows followed mainly the same protocol. Here only the cortex was aspirated until the alveus fibers above CA1 became visible. The silicon inset was shorter version (length 1.5 mm) of the one used for DG experiments.

### Two-photon calcium imaging

We used a commercially available two photon microscope (A1 MP, Nikon) equipped with a 25x long-working-distance, water-immersion objective (N.A.=1, WD=4 mm, XLPLN25XSVMP2, Olympus) controlled by NIS-Elements software (Nikon). GCaMP6s was excited at 940 nm using a Ti:Sapphire laser system (∼60 fs laser pulse width; Chameleon Vision-S, Coherent) and a fiber laser system at 1070 nm (55 fs laser pulse width, Fidelity-2, Coherent) to excite jRGECO1a (see **Supplementary Fig. 1B**). Emitted photons were collected using gated GaAsP photomultipliers (H11706-40, Hamamatsu). Movies were recorded using a resonant scanning system at a frame rate of 15 Hz and duration of 20 minutes per movie.

### Habituation and behaviour on the linear track

Experiments were performed in head fixed awake mice running on a linear track. Two weeks before the measurements, mice were habituated to the head fixation. Initially mice were placed on the treadmill without fixation for 5 minutes at a time. Subsequently, mice were head-fixed, but immediately removed if signs of fear or anxiety were observed. These habituation sessions lasted 5 minutes each and were carried out three times per day, flanked by 5 minutes of handling. During the following 3-5 days, sessions were extended to 10 minutes each. After habituation, mice ran well on the treadmill for average distances between 30 and 40 meters per session (see **Supplementary Fig. 1D**). The treadmill we implemented was a self-constructed linear horizontal treadmill, similar to (***Royer et al., 2012***)). The belt surface was equipped with tactile cues (see **Supplementary Fig. 1C**). Belt position and running speed were measured by modified optical mouse sensors. In addition, the pupil diameter was measured using a high-speed camera (Basler Pilot). All stimulation and acquisition processes were controlled by custom-made software written in LabView. Detailed construction plans and LabView software are available upon request.

### Data analysis, 2-photon imaging

All analysis on imaging data and treadmill behaviour data were conducted in MATLAB using standard toolboxes, open access toolboxes and custom written code. To remove motion artifacts, recorded movies were registered using a Lucas– Kanade model (***Greenberg and Kerr, 2009***). Individual cell fluorescence traces were identified and Ca^2+^ events were deconvolved using a constrained nonnegative matrix factorization based algorithm (***Pnevmatikakis et al., 2016***). All components were manually inspected and only those kept that showed shape and size of a granular cell and at least one significant Ca^2+^-event in their extracted fluorescence trace. We binarised individual cell fluorescence traces by converting the onsets of detected Ca^2+^ events to binary activity events. MPP input bulk signal was analyzed by setting a region of interest in the molecular layer. For that a threshold of 50 % maximum fluorescence was used within the field of view on the average projection of the movie. The bulk fluorescence signal trace was calculated as the average signal of the defined region of interest in each frame. The baseline for the bulk signal was defined as the low pass filtered signal of the raw trace with a cutoff frequency of 0.01 Hz using a Butterworth filter model. We did not observe any indication of epileptiform activity in Thy1-GCaMP6 (GP4.12Dkim/J) mice, as already reported (***Steinmetz et al., 2017***).

### Network activity

To define events of synchronized activity we first searched for Ca^2+^-event onsets occurring simultaneously within a moving time window of 200 ms (3 frames). We then defined the distribution of synchronous events that could arise by chance in each individual session. To achieve this, we created a null-distribution of synchronous events by randomly shuffling individual onset times a thousand times. We set the minimal threshold for network events in each individual session at a number of cells where less than 0.1% of events (p<0.001) could be explained by chance. Orthogonality between pairs of network events was assessed using cosine-similarity measures. Therefore, population vectors of all network events were multiplied using the normalized vector-product in a pair-wise manner. To test which fraction of orthogonal pairs could be explained by chance, we generated a null distribution by randomly reassigning the cell participations to different population vectors a 1000 times. We then tested the real fraction of orthogonal pairs against the fraction derived from the shuffled data.

### Spatial tuning

To address spatial tuning of activity in sparsely coding GCs we used spatial tuning vector analysis (***Danielson et al., 2016***). We restricted analysis to running epochs, where a running epoch was defined as an episode of movement with a minimal duration of 2.5 s above a threshold of 4 cm/s in a forward direction. Only cells with 4 or more event onsets during running epochs were included in the analysis. We calculated the mean of the vectors pointing in the mouse direction at the times of transient onsets, weighted by the time spent in that bin. We addressed statistical significance by creating the null distribution for every spatially tuned cell. This was achieved by randomly shuffling the onset times and recalculating the spatial tuning vector 1000 times. The p value was calculated as the percentage of vector lengths arising from the shuffled distribution that was larger than the actual vector length.

### Velocity tuning

To analyze speed modulated activity of GCs, velocity values were divided in 20 evenly sized bins between 0 and the maximum velocity of the animal. We calculated the mean □F/F at all times the animal was running at velocities within each specific velocity bin. Putative speed cells were those granule cells that showed a Pearson’s r of at least 0.9. To further exclude the possibility that correlations arise by chance we shifted the individual □F/F traces with respect to the behaviour randomly in the time domain a 1000 times. The cell was considered a significant speed coder if the shuffle-data r-values were below the original one in at least 95 % of the cases.

### Hierarchical cluster analysis

To find ensembles of correlated activity within network activity, we focused only on those granule cell Ca^2+^ events that occurred within network events. We calculated the correlation matrix using Pearson’s r for all cell combinations. To identify clusters of correlated cells we used agglomerative hierarchical cluster trees. Clusters were combined using a standardized Euclidean distance metric and a weighted average linkage method. Clusters were combined until the mean of the cluster internal r-value reached a significance threshold. To define the significance threshold we created a null-distribution of r-values from randomized data sets. Data was shuffled by randomly reassigning individual cell events to different network events a 1000 times. The 95 percentile of the created r-value null-distribution was used as the threshold for the clustering procedure. Only clusters in which the mean r-value exceeded the threshold obtained from the null distribution were considered for further analysis.

### Principal Component Analysis

To perform Principal Component Analysis (PCA) and Independent Component Analysis (ICA) we used standard MATLAB procedures and calculated the maximal number of components. Gaussian Process Factor Analysis (GPFA) was conducted using a formerly described procedure and toolbox (***Yu et al., 2009***), that we adapted for Ca^2+^-imaging data. Principals were calculated using singular value decomposition (SVD) of the data x of size *N* by *T*, where *N* is the number of cells, *T* the number of recorded frames and the rows of x are the z-scored Δ*F*/*F* traces, decomposing the data-matrix as

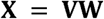

where V is an orthogonal matrix whos columns are the principal components, and W is a matrix of associated weights. For an analysis of population activity patterns relative to spatial location, we projected the animal position onto PCA trajectories, allowing us to identify loops in component space reflecting complete rounds on the belt. Further we projected all individual component amplitudes onto the position of the mouse to detect repetitive patterns. This analysis had comparable results for PCA, as well as ICA and GPFA (**Supplementary Fig. 5A-C** for dentate gyrus, **D-F** for CA1).

For further analysis we restricted the number of components so that 50 % of variance in each individual data set was explained. To compare running and network related epochs we calculated principal components V_r__un_ and V_net_ independently from each other so that

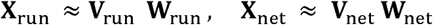

where x_ru__n_ contains all the data from epochs of running and x_net_ the data from 2 s windows around all network events. To calculate the similarity between these two bases, the covariance of x_run_ was projected into the principal space of the network activity

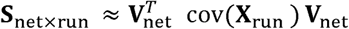

where S_net×run_ is the matrix of projected co-variances and trace(S_net×run_) quantifies the amount of projected co-variance.This number was normalized to the total amount of covariance of locomotion activity in the locomotion principal space trace(S_run×run_).

To compare our results against chance level, we used all traces recorded during immobility, shuffled those with respect to each other in the time domain. We used the original network times to create a random principal direction space V_rand_ and calculated the projected co-variances of x_run_ into the random-network space as trace(S_rand×run_) and repeated this procedure 1000 times. The p-value was calculated as the percentage of random projections that exceeded the initial value.

Additionally, we used two alternative approaches to quantify similarity between the PCA bases. First a similarity factor S_PCA_ as described by Krzanowski (***Krzanowski, 1979***).

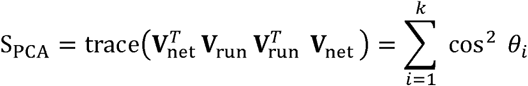

where *θ*_*i*_ is the angle between the *i*’th principal directions of V_run_ and V_net_. Further the Eros similarity factor as described in (***Yang, K., Shahabi, C., 2004***) was used:

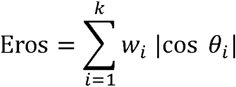

where w_*i*_ is a weighting factor. All measures delivered comparable results as compared to shuffled data.

### In-vitro patch-clamp experiments

Acute slices were prepared from mice expressing NpHR-eYFP in GCs. NpHR expression was induced by rAAV mediated gene transfer (rAAV2/1-DOI-eNpHR3-eYFP) into Prox1-Cre mice (see below for virus injection protocols). >2 weeks after virus injection, animals were deeply anesthetized with Isoflurane (Abbott Laboratories, Abbot Park, USA) and decapitated. The head was instantaneously submerged in ice-cold carbogen saturated artificial cerebrospinal fluid (containing in mM: NaCl, 60; sucrose, 100; KCl, 2.5; NaH_2_PO_4_, 1.25; NaHCO_3_, 26; CaCl_2_, 1; MgCl_2_, 5; glucose, 20) and the brain removed. Horizontal 350 µm thick sections were cut with a vibratome (VT1200 S, Leica, Wetzlar, Germany). Slices were incubated at 35 °C for 20 to 40 minutes and then stored in normal ACSF (containing in mM: NaCl, 125; KCl, 3.5; NaH_2_PO_4_, 1.25; NaHCO_3_, 26; CaCl_2_, 2.0; MgCl_2_, 2.0; glucose, 15) at room temperature. Recordings were performed in a submerged recording chamber at 33-35 °C under constant superfusion with carbogen saturated ACSF (3 ml/min). Visually identified GCs were recorded in current clamp using a low chloride intracellular solution (containing in mM: K-gluconate, 140; 4-(2-hydroxyethyl)-1-piperazineethanesulfonic acid (HEPES-acid), 5; ethylene glycol tetraacetic acid (EGTA), 0.16; MgCl_2_, 0.5; sodium phosphocreatine, 5) and a Multiclamp 700B and Digidata 1322A (Molecular Devices). Illumination (∼560nm, ∼1mW) was achieved through the Objective. Action potential frequencies were calculated using the smallest current injection yielding at least 4 action potentials.

### Light fiber implantation for behavioural experiments

Mice were injected with buprenorphine (0.05 mg/kg BW) 30 minutes before inducing anesthesia using 3.5 % isoflurane for induction and 1-1.5 % for maintenance. Mice were placed in a stereotactic frame (Kopf Instruments) and the scalp opened with surgical scissors after disinfecting it with iodine solution and applying local anesthetic (10% lidocaine). The skull was thoroughly cleaned using 2% H_2_O_2_, covered with a thin layer of two-component dental adhesive (Optibond) and the surrounding wound sealed with tissue glue (Vetbond). Small craniotomies were performed above the target sites and 500 nl virus suspension (rAAV2/1-Ef1a-DOI-NLS-eYFP for controls, rAAV2/1-EF1a-DOI-eNpHR3-eYFP for experimental group) was bilaterally injected using a 34 G syringe (Nanofill Syringe, World Precision Instruments, Inc.) at a speed of 50 nl/min. After each injection, the syringe was kept in place for 5 minutes to ensure permeation of the virus into the parenchyma. Coordinates for viral injections into dorsal dentate gyrus were: (from bregma): AP: −2.3; ML: −/+ 1.6 and DV: 2.5 mm). Afterwards, fiber optic cannulae of 200 µm diameter (NA: 0.39, CFMLC12, Thorlabs) were bilaterally implanted at (from bregma): AP: −1.7; ML: −/+ 1.35 and DV: 1.7 and fixed to the skull with a layer of flowable opaque composite (Tetric Evoflow) topped by multiple layers of dental cement (Paladur, Heraeus). Finally, antibiotic cream (Refobacin, Almirall) was applied to the wound and the animals received ketoprofen (5 mg/kg BW) s. c. Analgesia was applied post-surgery by injecting ketoprofen (5 mg/kg BW) s. c. after 24, 48, and 72 hours. All mice recovered for at least 3 weeks after surgery before the start of behavioural experiments.

### Behavioral experiments

To test the effect of optogenetic inhibition of granule cells, 18 heterozygous Prox1-Cre animals (3 male, 15 female) between the age of 6 and 9 months were used. Males were single caged and females were group-caged (2-4 individuals per cage) in standard mouse cages under an inverted light-dark cycle with lights on at 8 pm and ad libitum access to food and water. Prior to experiments, animals were randomly assigned to the control or experimental group. All experiments were conducted during the dark phase of the animals. Prior to experiments, mice were handled by the experimenter for at least 5 days (at least 5 min/day). On experimental days, animals were transported in their home cages from the holding facility to the experimental room and left to habituate for at least 45 minutes. All experiments were performed under dim light conditions of around 20 Lux. Animals were also habituated to the procedure of photostimulation by attaching a dual patch cord for around 10 minutes to the bilaterally implanted light fibers and letting them run in their home cage on multiple days.

For optogenetic stimulation, 561 nm laser light (OBIS/LS FP, Coherent, Santa Clara, CA) was delivered bilaterally into the implanted optical fibers using a dual patch cord (NA: 0.37, Doric lenses, Quebec, Canada) and a rotary joint (FRJ, Doric lenses, Quebec, Canada) located above the behavioural test arena. The laser power was set to 5 mW at the tip of the fiber probes.

We used a pulsed laser light at 20 Hz with a 50% duty cycle instead of continuous illumination to keep the light-induced heat effect minimal in the brain. Previous studies have shown that continuous light delivery to brain tissues can cause a temperature increase of up to 2°C (***Owen et al., 2019***). It has been reported that changes in temperature can alter neuronal physiology of rodents (***Stujenske et al., 2015***) and birds (***Long and Fee, 2008***). We simulated the light-induced heat effect using the model developed by (***Stujenske et al., 2015***). We first examined the continuous light photostimulation of 561 nm light with 5 mW output power. The temperature increase reached a steady-state and was found to be 0.7°C in 60 seconds. For the pulsed light stimulation used in this study, the temperature change did not exceed 0.3°C in 60 seconds (see **Supplementary Fig. 7K-M**). Both pulsed and continuous stimulation resulted in efficient silencing of granule cell activity (**Supplementary Fig. 7E-J**). The average illumination times were not significantly different between eNpHR and eYFP control groups in any of the experiments. To apply photostimulation only during immobility, we used a closed loop system employing EthoVision 8.5 that opened an optical shutter (SH05, Thorlabs) in the light path of the laser via a TTL pulse only when the tracking software detected that the body center of the animal was moving less than 5 cm/s and closed the shutter if the speed exceeded 5 cm/s over a period of 0.5 seconds. Behaviour was recorded using EthoVision 8.5 software (Noldus, Netherlands) and an IR camera with a frame rate of 24 Hz. Videos were stored on a computer for offline analysis.

### Object pattern separation task

We used the object location memory test as a test for spatial learning ability. We used a circular arena (diameter: 45 cm) made of red Plexiglas with 40 cm high walls as described in (***van Goethem et al., 2018***). Two pairs of almost identical building blocks (4 × 3 × 7 cm) made of either plastic or metal were used as objects. All building blocks were topped with a little plastic cone to prevent animals from climbing onto the objects. To habituate animals to the arena and the experimental procedure, they were taken out of their home cage and placed into an empty cage only with bedding for five minutes in order to increase their exploration time. After connecting the mice to the photostimulation apparatus via the light fiber, they were placed into the empty arena for 10 minutes with no laser light. On test days, mice were again first placed into a new empty cage for 5 minutes, then connected to the photostimulation apparatus and placed into the arena for 5 minutes. The arena contained two identical objects placed in the middle of the arena with an inter-object distance of 18 cm. The animals were allowed to freely explore the arena and the objects. After 5 minutes, the animals were transferred into their home cage for 85 minutes. For the recall trial, everything was done identically to the previous acquisition trial, but one of the two previously encountered objects was displaced. Experiments for individual displacement configuration were repeated in some cases up to three times with an interval of two days and the discrimination indices averaged. The following variants of this task regarding object displacement were performed. In a first set of experiments, variable displacements (3, 6, 9 and 12 cm) were used, with inhibition of granule cells carried out during the entire acquisition and recall trial (schematic in **Supplementary Fig. 7A-C**). In a second set of experiments a fixed, intermediate degree of displacement (9 cm, position 4 in **Supplementary Fig. 7C**) was used, and inhibition of granule cells was carried out only during quiet immobility using a closed-loop system (see above) in either only the acquisition trials (**Fig. 5**) or only the recall trials (**Supplementary Fig. 8**). In a final set of experiments, we applied photostimulation in the acquisition trials only when the mouse was in quiet immobility and the nose point of the mouse at least 4 cm away from the objects (**Supplementary Fig. 8**). After each mouse, the arena and the objects were cleaned with 70 % ethanol. An experienced observer blinded to the experimental group of the animals manually scored the time the animals explored the displaced and the non-displaced object by sniffing with the snout in very close proximity to the objects and their head oriented towards them. We calculated the discrimination index based on the following formula: (time exploring displaced object – time exploring the non-displaced object) / (time exploring displaced object + time exploring the non-displaced object). Trials in which the total exploration time in an acquisition or recall trial was lower than 4 seconds were excluded from further analysis. Trajectory maps and occupancy plots were generated using custom-written MATLAB Scripts.

### Histology and microscopy

To verify successful viral transduction and window position, animals were deeply anesthetized with ketamine (80 mg/kg body weight) and xylazine (15 mg/kg body weight). After confirming a sufficient depth of anesthesia, mice were heart-perfused with cold phosphate buffered saline (PBS) followed by 4% paraformaldehyde (PFA) in PBS. Animals were decapitated and the brain removed and stored in 4% PFA in PBS solution. Fifty to 70 µm thick coronal slices of the hippocampus were cut on a vibratome (Leica). For nuclear staining, brain slices were kept for 10 min in a 1:1000 DAPI solution at room temperature. Brain slices were mounted and the red, green and blue channel successively imaged under an epi fluorescence or spinning disc microscope (Visitron VisiScope). In optogenetic inhibition experiments, post-hoc microscopy was used to confirm successful expression of NpHR-eYFP. Of the animals used, 3 control animals showed no eYFP expression and one experimental animal lacked NpHR-eYFP expression. The animal lacking NpHR expression was excluded from the study. Control animals lacking eYFP expression were assumed to lack modulation of granule cell activity by illumination and were pooled with eYFP expressing control animals.

## Acknowledgements

We are very grateful to David Greenberg, Jason Kerr, and Damian Wallace for technical help and advice, as well as the supply of analysis algorithms. We gratefully acknowledge the support of Jonathan Ewell in editing the manuscript. We acknowledge the support of the Imaging Core Facility of the Bonn Technology Campus Life Sciences. The work was supported by the SFB 1089, Project C04 to HB, the Research Group FOR2715, the Research Priority Program SPP Computational Connectomics and EXC 2151 under Germanys Excellence Strategy of the Deutsche Forschungsgemeinschaft (DFG, German Research Foundation) to HB, and support of the Humboldt Foundation PSI to KG. ANH was supported by the IMPRS Brain and Behaviour.

