## Supplementary figures for "Dentate gyrus population activity during immobility supports formation of precise memories"

**Supplementary Figures 1-9:**

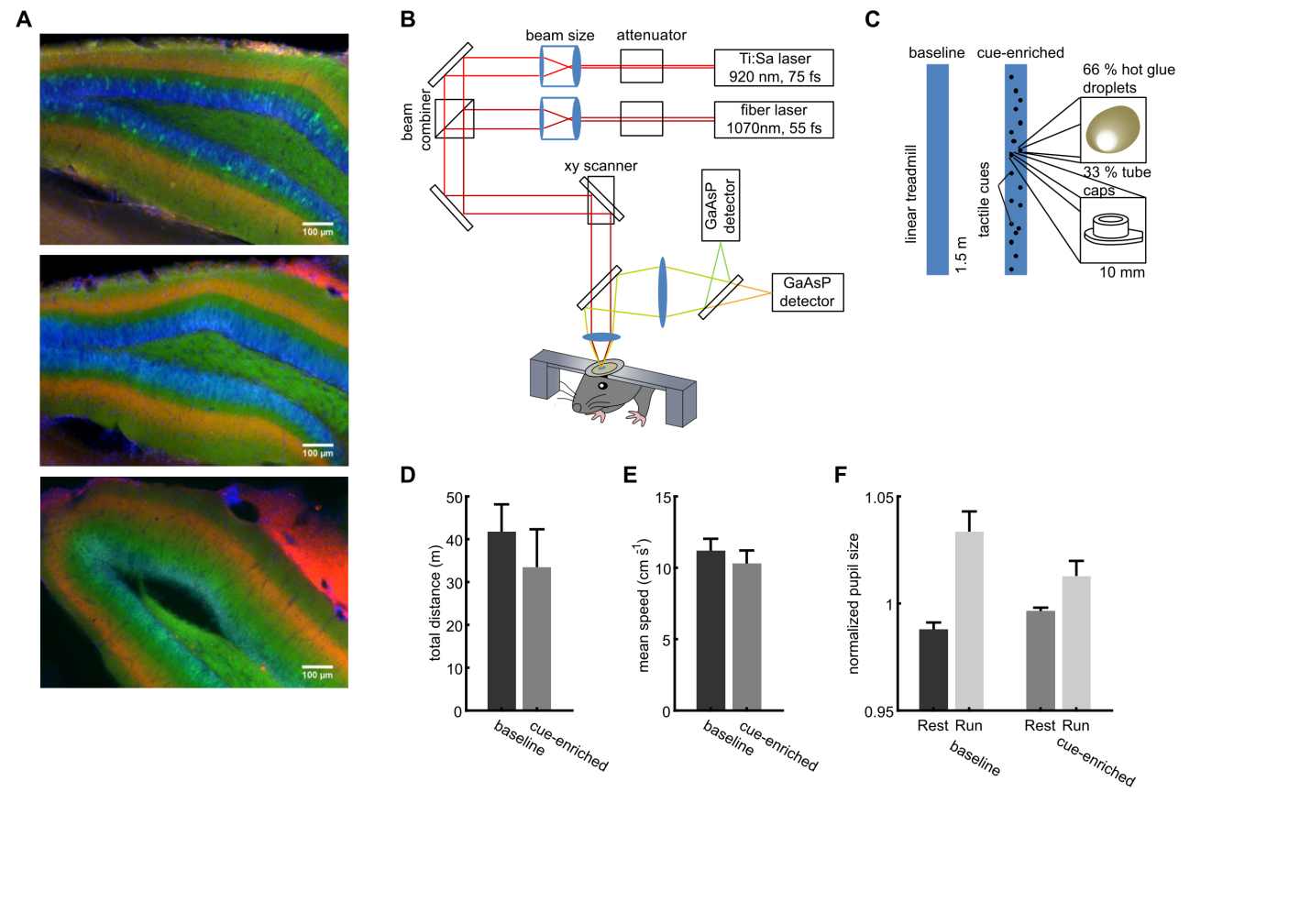

**Supplementary Fig. 1:** **Dual color two-photon imaging in the dentate gyrus.** **A,** Representative hippocampal sections from three different mice with expression of jRGECO in the MPP (red), GCaMP6s in granule cells (green), and DAPI as nuclear staining. **B,** Setup of the 2-photon microscope for dual-color two-photon imaging. To allow efficient excitation of both genetically encoded Ca^2+^ indicators, we established excitation with two pulsed laser sources at 920 and 1070 nm. **C,** Properties of the linear track and dimensions of the spatial cues. **D, E,** Locomotion on the linear track. Neither the total distance run on the linear track (d), nor the average running speed (e) differed significantly between the baseline and cue-enriched conditions (n.s., ANOVA, p=0.53 and 0.58, respectively). **F,** Measurement of pupil size under the different locomotor and cue conditions. Two-way ANOVA revealed significantly larger pupil sizes in running vs. resting mice (F_(1,6)_=17.08, p=0.0004). No differences between baseline and cue-enriched conditions were detected (F_(1,6)_=0.86, p=0.36). Linked to Fig. 1.

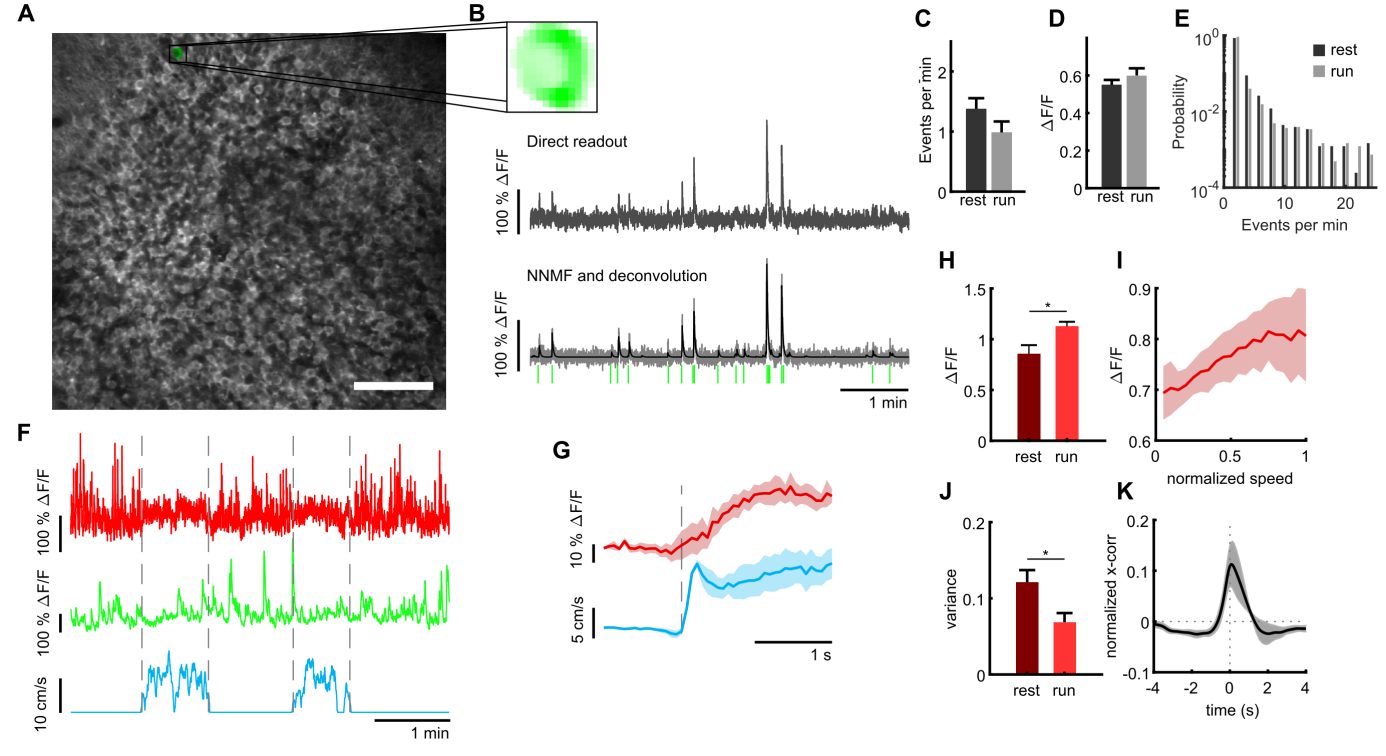

**Supplementary Fig. 2: Granule cell and MPP activity during locomotion on empty textured belt.** **A,** Image plane for granule cell recording in the dentate gyrus. **B,** One representative component of non-negative matrix factorization. Upper panel: spatial component as extracted from image stack. Middle panel: Corresponding ΔF/F trace generated from a ROI drawn around the corresponding somatic region. Lower panel: Extracted ΔF/F trace (gray) with deconvoluted trace (black). Identefied event onsets are depicted with vertical green lines. **C, D,** Frequencies of Ca^2+^ events (C) and magnitude of Ca^2+^ transients during each recording session (D) recorded during quiet immobility (rest, dark bars) and locomotion (run, light bars) **E**, Distribution of event frequencies for all cells. Black bars represent events during resting periods, gray bars represent frequencies during locomotion. **F,** Mean MPP activity (red) and the sum of all granule cell activities (green) for a representative section of a recording session. Dashed lines mark transition between resting and running periods (see blue line indicating running speed). **G,** Average MPP fluorescence and running speeds, both aligned to running onsets (dashed line). Shaded areas indicate standard error (n=4 animals). **H,** Mean fluorescence averaged during resting (dark red) and running (light red, n=4). **I,** Correlation of mean activity of the MPP input with the mouse running speed (n=4). **J,** Variance of MMP bulk signal during resting (dark red) and running (light red, n=4). **K,** Cross correlation of MPP bulk signal and summed GC signal during resting. Linked to Figure 1.

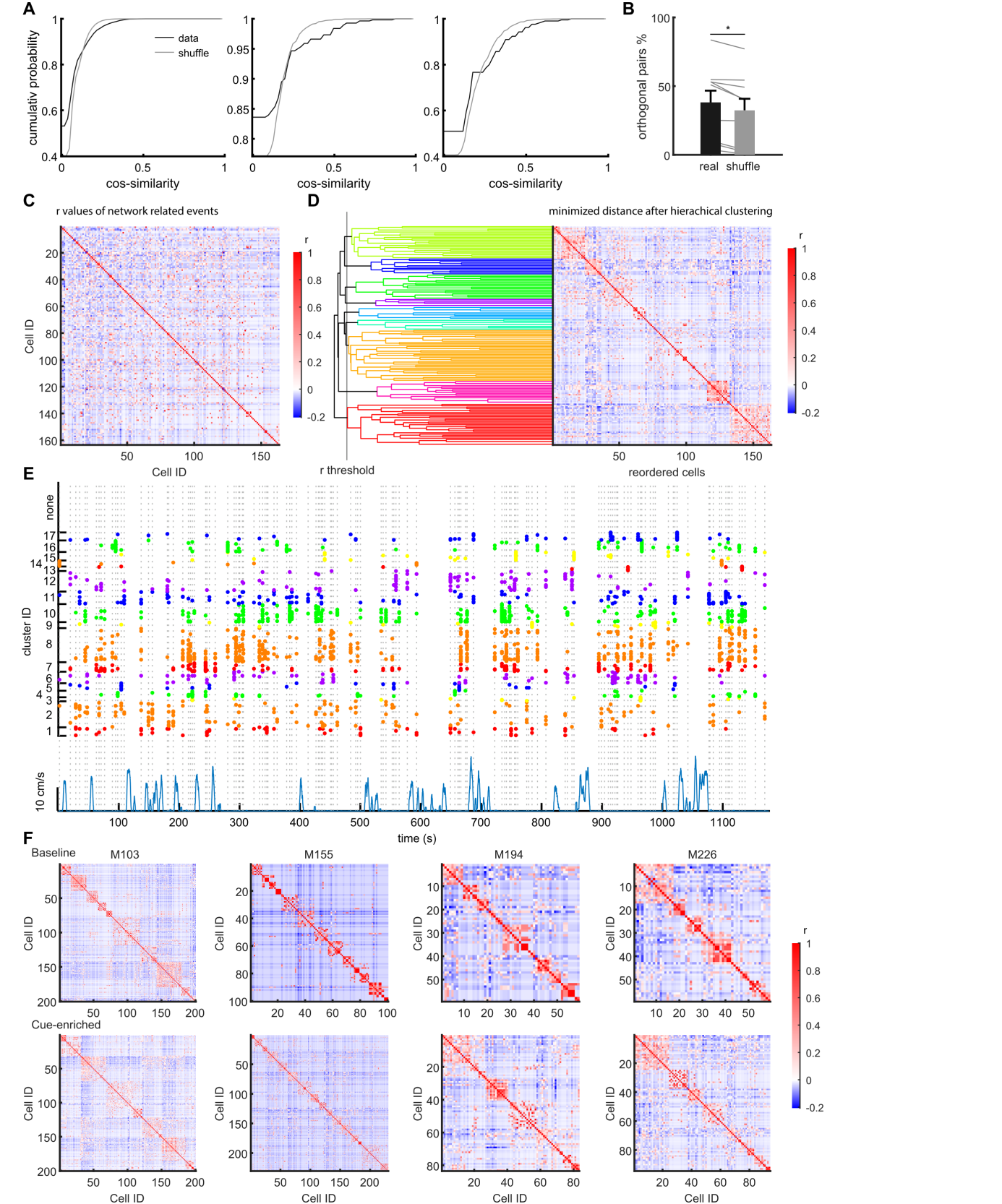

**Supplementary Fig. 3:** **Clustering of cells active during network events into correlated assemblies via a correlation matrix.** **A,** Representative distributions from three representative sessions for the cumulative probability to find similar vs. orthogonal pairs of network events using cosine similarity. Black line represents comparisons for real data pairs whereas in gray the mean from randomly shuffled data (see Methods). **B,** Percentage of orthogonal pairs using the zero values from panel A. The real data comparisons (black bar) contain significantly more orthogonal pairs compared to shuffled data (grey bar) **C,** Graphical representation of the correlation matrix using Pearson’s r for all cell combinations, with values for r being color coded. Data from one representative session in an individual mouse. **D,** Identification of clusters of correlated cells using agglomerative hierarchical cluster trees. Clusters were combined using a standardized Euclidean distance metric and a weighted average linkage method. Clusters were combined until the mean of the cluster internal r-value reached a significance threshold. The significance threshold was defined by creating a null-distribution of r-values from randomized data sets, and is indicated for this particular experiment with a vertical line. Right panel in d depicts the reordered correlation matrix showing clusters of highly correlated cells. **E,** Raster plot showing the occurrence of clusters identified in d over multiple episodes of running and immobility, showing the reactivation of clusters over time. Individual dots indicate participation of individual cells. Clusters are color-coded. Network events are indicated by vertical dashed lines. Running episodes are indicated at the lower border with the running speed (blue). Linked to Fig. 1. **F**, Examples of correlation matrices after hierarchical clustering from four mice. Upper row shows baseline condition and lower row cue-enriched condition.

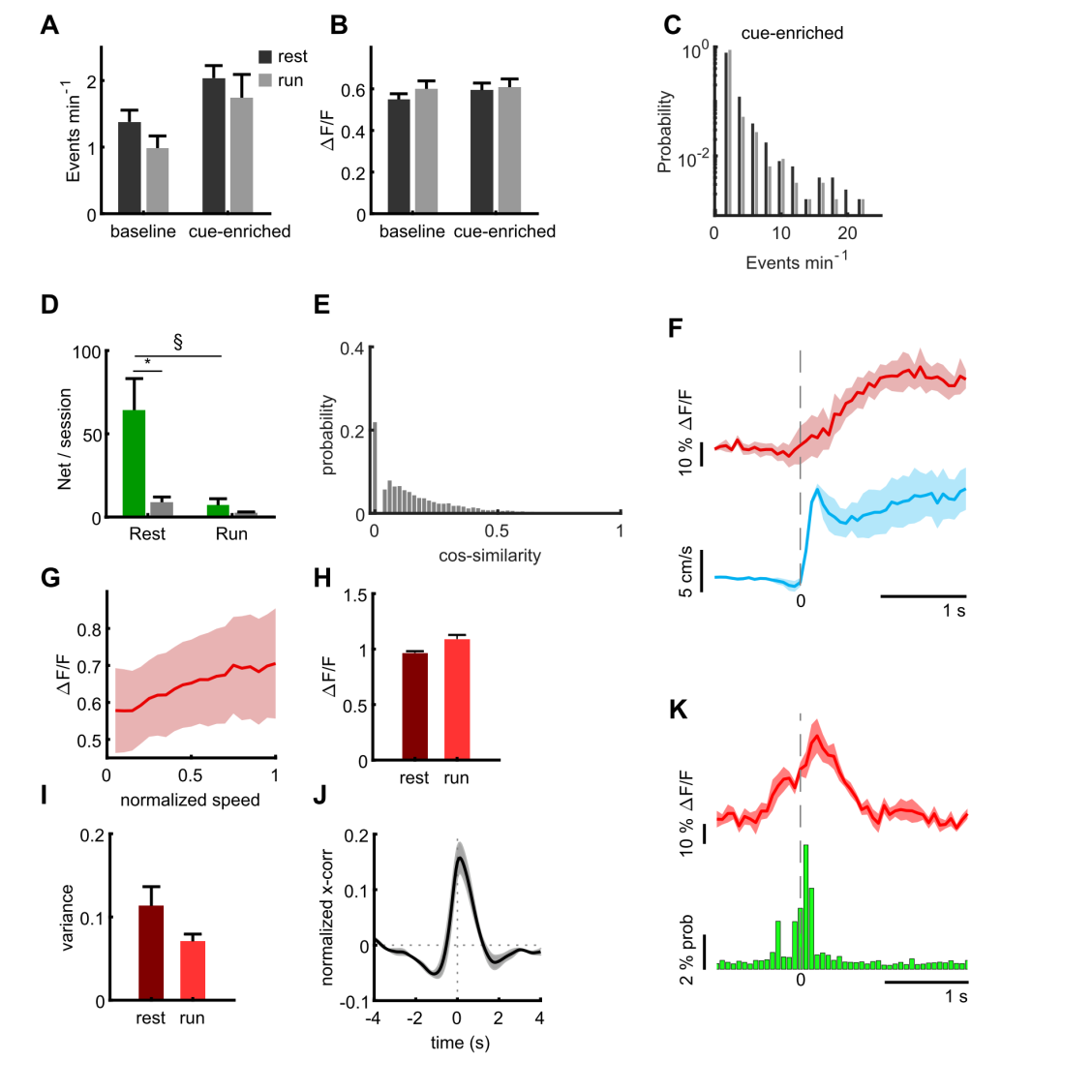

**Supplementary Fig. 4: Activity of granule cells and MPP inputs in cue-enriched conditions. A, B,** Average frequencies of Ca^2+^ events (A) and magnitude of Ca^2+^ transients (B) for baseline and cue-enriched conditions recorded during quiet immobility (rest, dark bars) and running (run, light bars). **C**, Distribution of event frequencies for all cells for the cue-enriched condition. **D,** Under cue-enriched conditions, network events also occurred mainly during immobility. Mean number of network events per recording session during running (light green) and resting (dark green). Grey bars depict shuffled data for each condition. ANOVA F_(3,28)_=8.6, p=0.0003, Bonferroni post-test resting vs. shuffled p=0.0019 indicated with asterisk, running vs. shuffled p=0.96). **E,** Similarity between network events under cue-enriched conditions. Similarity of population vectors computed for individual network events. Comparisons were carried out between all possible pairwise combinations of vectors and quantified using cosine similarity. **F-K,** Activity of MPP inputs in the cue-enriched condition. Shaded areas indicate standard errors. **F,** In cue-enriched conditions, MPP activity also increases at transitions from immobility to running (red line, MPP activity, blue line indicates running speed, n=4). **G,** Correlation of mean activity of the MPP input with the mouse running speed in a representative mouse in a cue-enriched trial (correlation coefficient for all mice: r=0.98, p=3.9*10^-13^). **H,** Mean fluorescence averaged during resting (dark red) and running (light red, n=4) in cue-enriched trials. ANOVA for running vs. immobile states F_(1,3)_=9.64, p=0.02. **I,** Variance of MPP bulk signal during resting (dark red) and running (light red, n=4) in cue-enriched trials. ANOVA for running fs. Immobile states F_(1,3)_=3.00, p=0.13. **J,** Cross correlation of MPP bulk signal and summed GC signal during resting. **K,** Average MPP activity (red) and probability of granule cells being active in cue-enriched trials, both aligned to the time point of network events. Linked to Fig. 3.

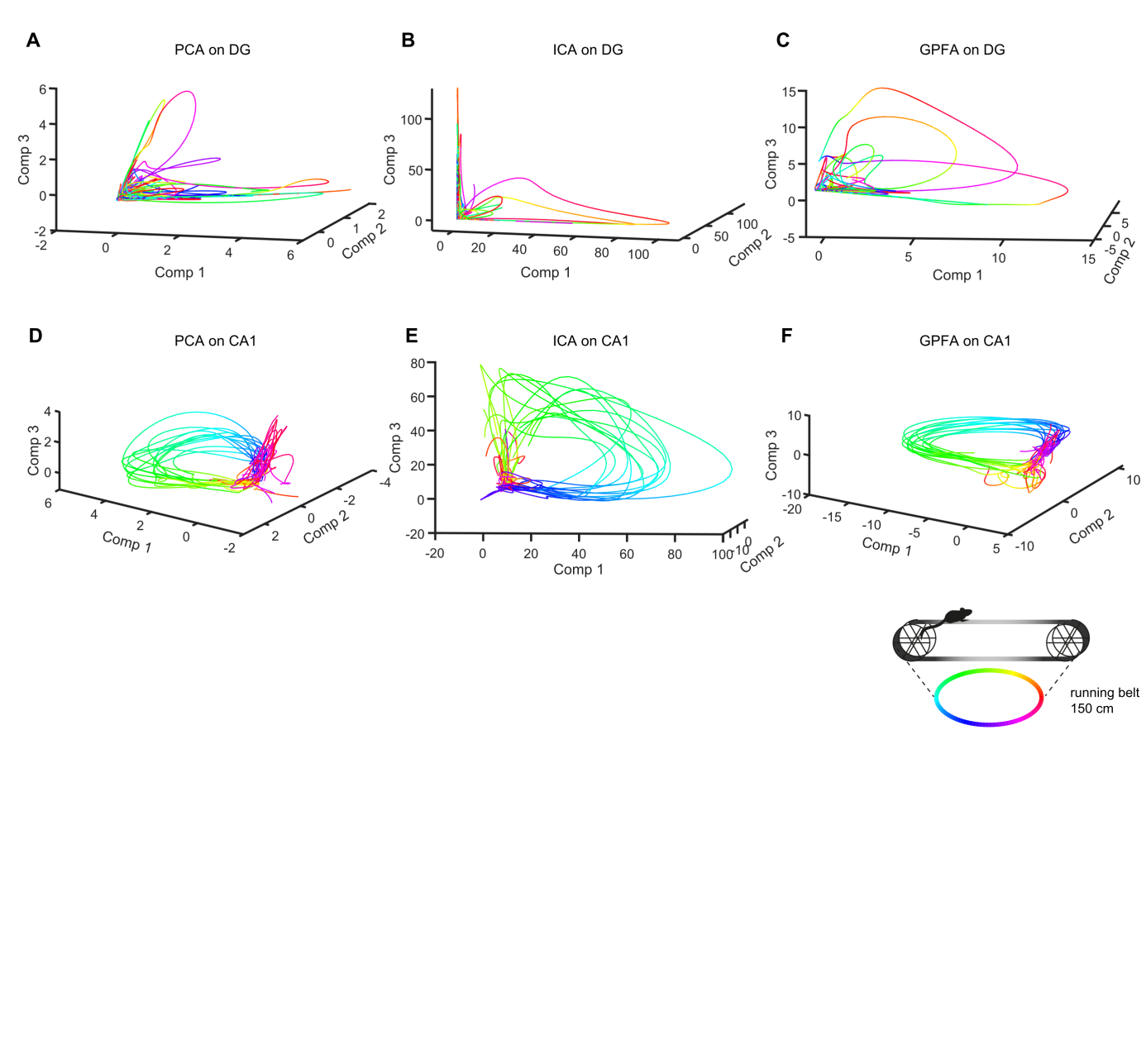

**Supplementary Fig. 5**: **Analysis of population activity in dentate gyrus and the CA1 subfield of the hippocampus using PCA, ICA and GPFA.** **A-C,** Upper panels depict the first three components from representative sessions (A: PCA, B: ICA, C: GPFA) plotted in a coordinate system. The color code refers to the place on the linear track, with the same locations represented in the same color. **D-F,** As in A-C, but for CA1 neurons. Note the smooth and repetitive trajectories. Linked to Fig. 4.

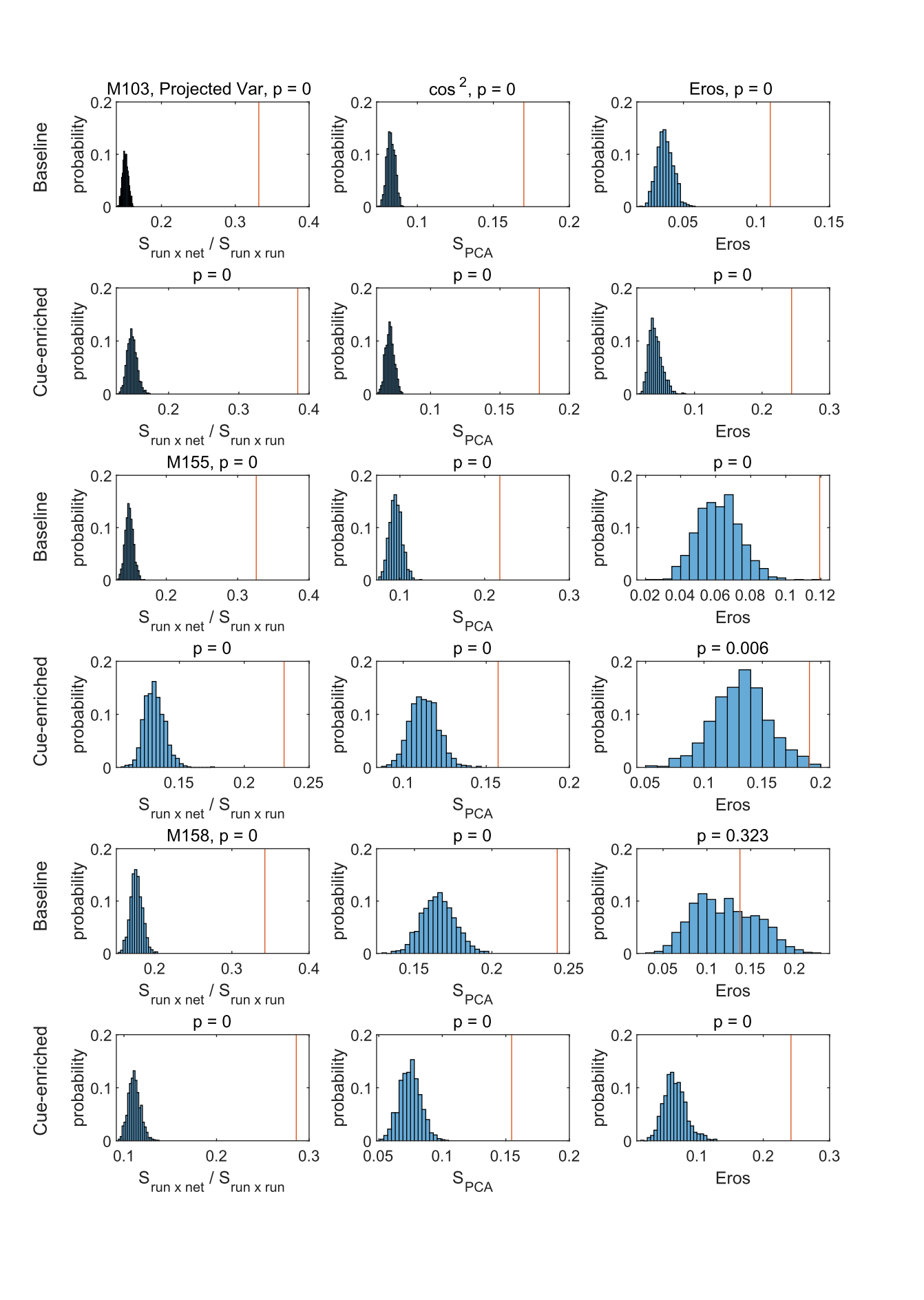

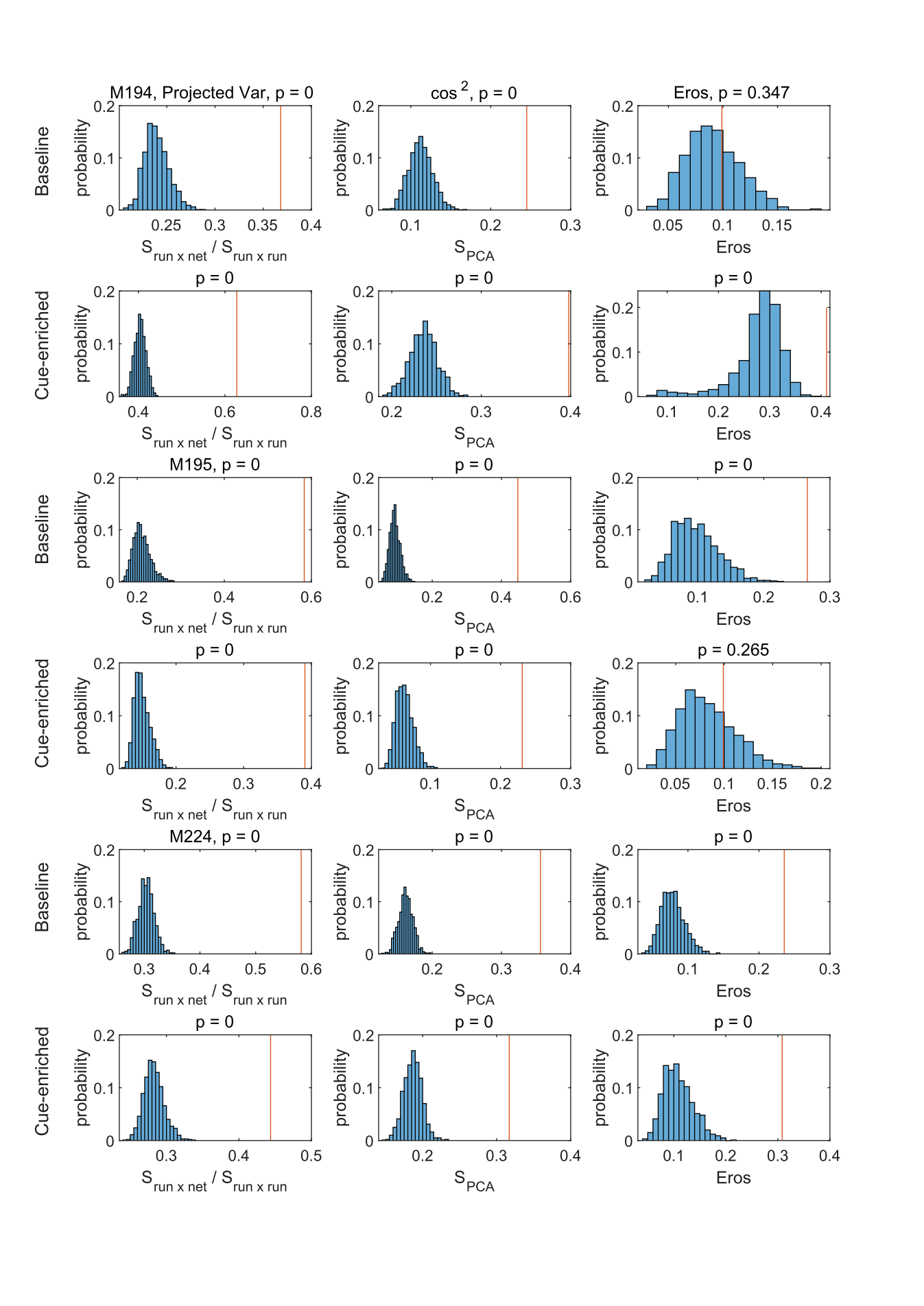

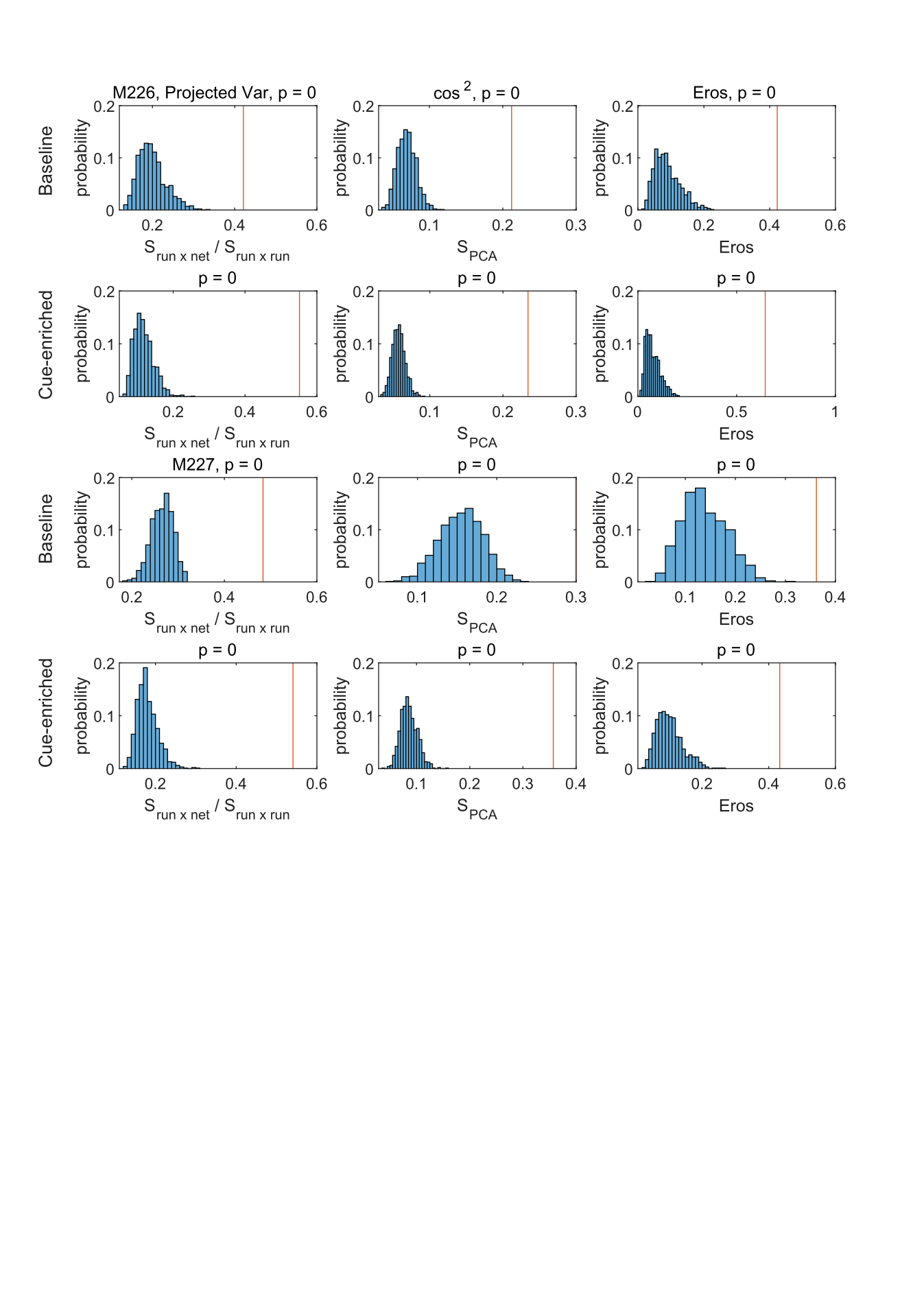

**Supplementary Fig. 6: Similarity of individual network events to population activity during running.** Data from all sessions under baseline and cue-enriched conditions are depicted (as indicated on the leftmost border of the figure) for all three measures. PCA similarity using the vector projection method introduced in this paper (see Methods), cosine similarity measures (***Krzanowski, 1979***) and EROS (***Yang, K., Shahabi, C., 2004***) are depicted in the leftmost, middle and rightmost columns, respectively. In all graphs, shuffled data distributions are shown in light blue, a vertical red line indicates the similarity value between network and locomotion related population activity in the particular session. P-values are indicated above each graph. In the variance projection method, values were normalized to the locomotion related variances projected into the locomotor states. As expected, this results in a high proportion of explained variance. Linked to Fig. 4.

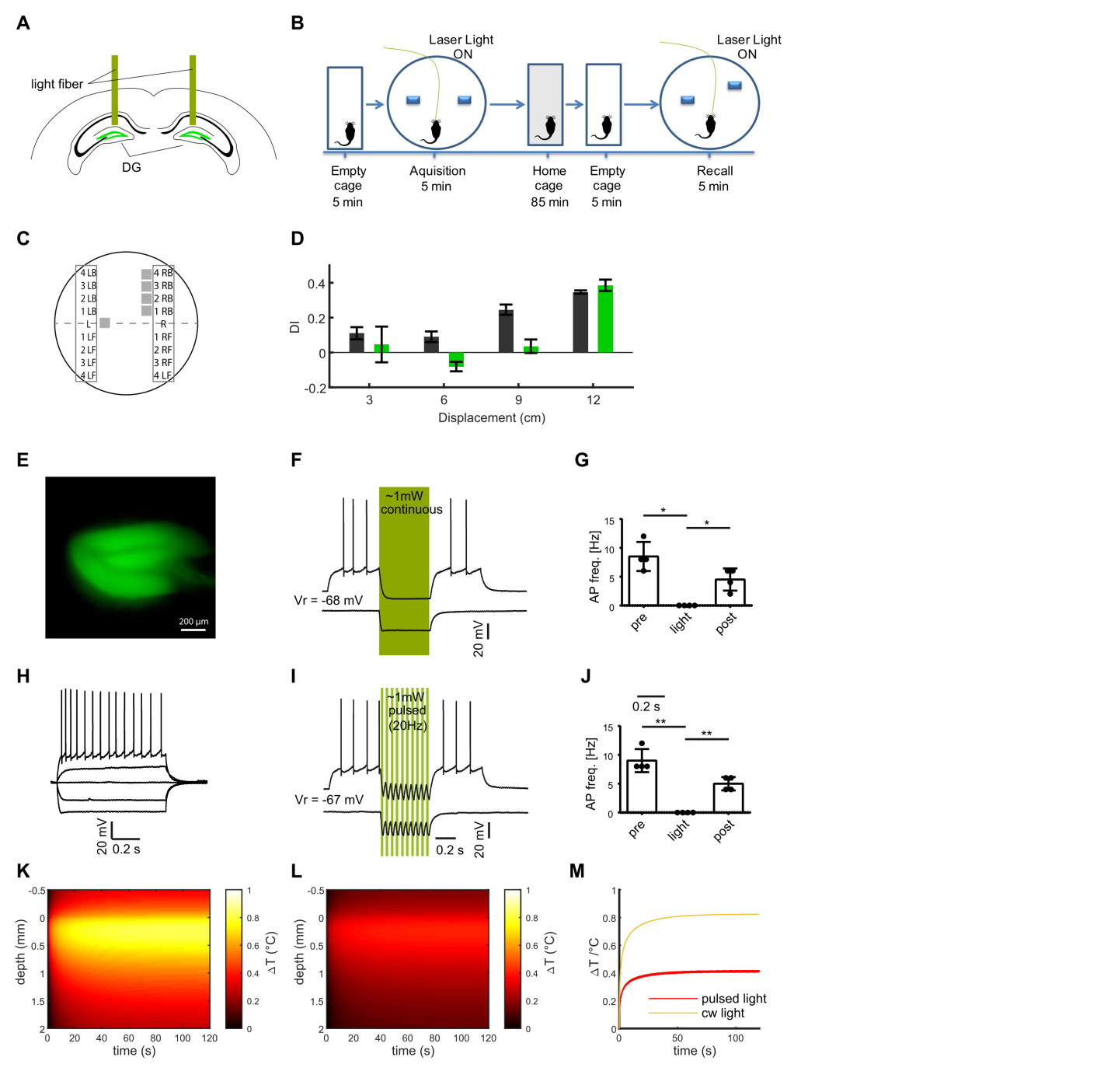

**Supplementary Fig. 7: Establishing a dentate gyrus-dependent variant of the object pattern separation task. A,** Schematic of the bilateral optogenetic inhibition of the dentate gyrus granule cells expressing eNpHR. **B,** Description of the task. Following familiarization with the object location, one of the objects is moved in a subsequent recall session, and the extent to which mice explore the moved vs. stationary object is examined. These trials can be repeated allowing to explore the effects of variable movement of the objects. **C,** Schematic of possible object locations for the displaced object. Displacement was randomized for each animal, such that either the left or the right object was displaced, in either a forward or back direction (LB i.e. corresponding to left, back, and RF to right, forward). The experiment used four possible new locations along a vertical axis, increasing from minor displacement (3 cm) to maximal displacement, indicated by numbers 2-5. **D,** Results of light-based inhibition of granule cells during acquisition and recall trials for different degrees of object separation indicate on the x-axis (eNpHR group, n=4, green bars) vs. an eYFP expressing control group (n=3, black bars). The effect of granule cell inhibition is most pronounced for intermediate degrees of object movement. **E,** Wide-field image of a hippocampal slice showing expression of NpHR-eYFP in granule cells. **F,** inhibition of granule cell firing evoked with long current injections by light-based activation of NpHR (yellow vertical bars) with continuous stimulation. **G,** quantification of firing rates before, during and after illumination for continuous stimulation. **H,** Representative, typical discharge behaviour of a granule cell. **I,** like F with pulsed stimulation at 50% duty cycle and 20 Hz (d). **J,** like G with pulsed stimulation. **K, L,** Estimation of light-induced warming within brain tissue for continuous illumination (**K**) or pulsed illumination at 50% duty cycle and 20 Hz (**L**) at intensities used for the behavioural experiments. Predicted temperature changes are plotted as a function of time and depth. **M,** Analysis of warming over time, showing that the warming effects of pulsed light stimulation are asymptotic, and remain below 0.4 °C at a distance of 300 µm from the fiber front end. Linked to Fig. 5.

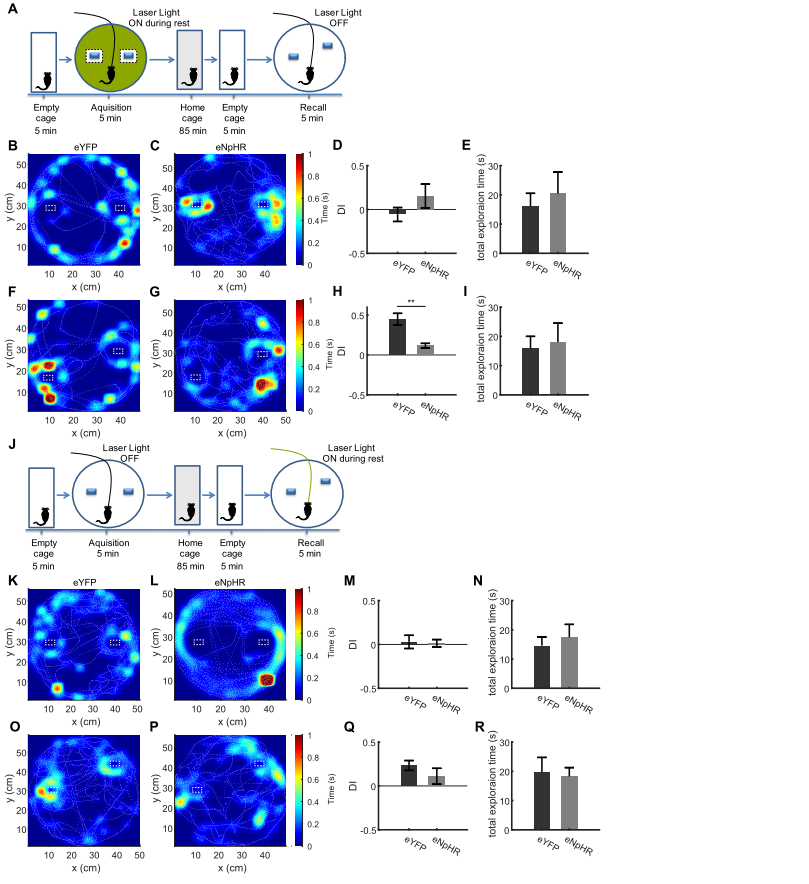

**Supplementary Fig. 8: Inhibition of dentate granule cell activity during immobility in the acquisition trial only in non-object locations impairs memory formation in the OPS task. A,** Description of the experimental protocol. In the acquisition phase, mice were familiarized with an arena containing two objects. Following an intermediate period of 90 minutes, the mice were placed in the same arena in which one object was moved slightly. Inhibition of granule cells was carried out only during periods of quiet immobility in the acquisition trial, and only if the periods of immobility were not adjacent to the objects. **B, C,** Representative sessions from acquisition trials in control (eYFP) mice (b, n=10) and mice expressing eNpHR in granule cells (c, n=4) showing the tracking data (dashed white lines), as well as occupancy within the open field as a heat map. **D,** Discrimination index quantifying relative exploration times of the two objects, with 0 values indicating equal exploration (see Methods). Comparison between groups n.s., t-test with Welch correction p=0,3537. **E,** Total time spent exploring the objects in the eYFP and eNpHR groups during the acquisition trial. Comparison between groups n.s., t-test with Welch correction p=0,6126. **F, G,** Representative sessions from recall sessions. **H,** Discrimination index for recall trials, showing a significant reduction in the recognition of the displaced object in the eNpHR group. t-test with Welch correction p=0,0018. **I,** Total time spent exploring the objects in the eYFP and eNpHR groups during the recall trial. Comparison between groups n.s., t-test with Welch correction p=0,4285. **J,** Inhibition of dentate granule cell activity during immobility does not interfere with recall in the OPS task.. In the acquisition phase, mice were familiarized with an arena containing two objects. Following an intermediate period of 90 minutes, the mice were placed in the same arena in which one object was moved slightly. Inhibition of granule cells was carried out only during periods of quiet immobility in the recall trial. **K, L,** Representative sessions from acquisition trials in control (eYFP) mice (b, n=11) and mice expressing eNpHR in granule cells (c, n=6) showing the tracking data (dashed white lines), as well as occupancy within the open field as a heat map. **M,** Discrimination index quantifying relative exploration times of the two objects, with 0 values indicating equal exploration (see Methods). Comparison between groups n.s., t-test with Welch correction p=0.8613. **N,** Total time spent exploring the objects in the eYFP and eNpHR groups during the acquisition trial. Comparison between groups n.s., t-test with Welch correction p=0.6097. **O, P,** Representative sessions from recall sessions. **Q,** Discrimination index for recall trials, showing no significant reduction in the recognition of the displaced object in the eNpHR group. Comparison between groups n.s., t-test with Welch correction p=0.2802. **R,** Total time spent exploring the objects in the eYFP and eNpHR groups during the recall trial. Comparison between groups n.s., t-test with Welch correction p=0.8236. Linked to Fig. 5.

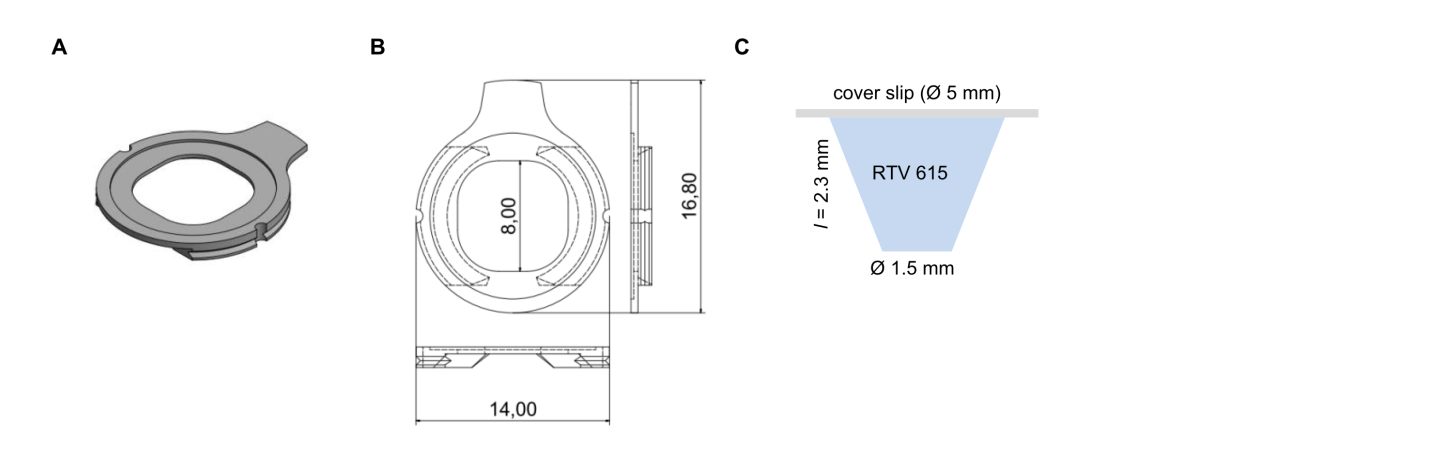

**Supplementary Fig. 9: Details of head fixation.** **A, B,** Dimensions of flat, custom head fixation ring. **C,** conical transparent inset used to maximize NA in deep imaging. Linked to Methods.

**Supplementary Movies:**

**Supplementary Movie 1:** Movie showing activity of granule cells and MPP, corresponding to Fig. 1a, b.

**Supplementary Movie 2:** Movie showing network events, corresponding to Fig. 2a
